# Identification and drug-induced reversion of molecular signatures of Alzheimer’s disease onset and progression in *App^NL-G-F^*, *App^NL-F^* and 3xTg-AD mouse models

**DOI:** 10.1101/2021.03.17.435753

**Authors:** Eduardo Pauls, Sergi Bayod, Lídia Mateo, Víctor Alcalde, Teresa Juan-Blanco, Takaomi C Saido, Takashi Saito, Antoni Berrenguer-Llergo, Camille Stephan-Otto Attolini, Marina Gay, Eliandre de Oliveira, Miquel Duran-Frigola, Patrick Aloy

**Author notes:** These authors have contributed equally to this work (Eduardo Pauls, Sergi Bayod, Lídia Mateo).

## Abstract

Alzheimer’s disease (AD) is the most common form of dementia. Over fifty years of intense research have revealed many key elements of the biology of this neurodegenerative disorder. However, our understanding of the molecular bases of the disease is still incomplete, and the medical treatments available for AD are mainly symptomatic and hardly effective. Indeed, the robustness of biological systems has revealed that the modulation of a single target is unlikely to yield the desired outcome and we should therefore move from gene-centric to systemic therapeutic strategies. Here we present the complete characterization of three murine models of AD at different stages of the disease (i.e. onset, progression and advanced). To identify genotype-to-phenotype relationships, we combine the cognitive assessment of these mice with histological analyses and full transcriptional and protein quantification profiling of the hippocampus. Comparison of the gene and protein expression trends observed in AD progression and physiological aging revealed certain commonalities, such as the upregulation of microglial and inflammation markers. However, although AD models show accelerated aging, other factors specifically associated with Aβ pathology are involved. Despite the clear correlation between mRNA and protein levels of the dysregulated genes, we discovered a few proteins whose abundance increases with AD progression, while the corresponding transcript levels remain stable. Indeed, we show that at least two of these proteins, namely lfit3 and Syt11, co-localize with Aβ plaques in the brain. Finally, we derived specific Aβ-related molecular AD signatures and looked for drugs able to globally revert them. We found two NSAIDs (dexketoprofen and etodolac) and two anti-hypertensives (penbutolol and bendroflumethiazide) that overturn the cognitive impairment in AD mice while reducing Aβ plaques in the hippocampus and partially restoring the physiological levels of AD signature genes to wild-type levels.

**Teaser:** The comprehensive characterization of three AD mouse models reveals disease signatures that we used to identify approved drugs able to modify the etiology of the pathology and overturn cognitive impairment.

## Introduction

Alzheimer’s disease (AD) is the most common form of dementia. The accumulation of amyloid-beta (Aβ) peptide in the form of plaques and the formation of intracellular Tau neurofibrillary tangles in the brain are the main pathological hallmarks of this neurodegenerative disease (*1*). Mutations in genes that are part of the Aβ processing pathway (e.g. *APP*, *PSEN1* and *PSEN2*) cause infrequent cases of hereditary AD (*2*). This observation thus reinforces the hypothesis that Aβ accumulation plays a necessary role in AD onset. At a later stage, Aβ aggregation induces a series of molecular changes that lead to Tau hyper-phosphorylation and intracellular fibrillation, which in turn causes neuronal death and neurodegeneration (*3*).

Transgenic mice that develop extensive Aβ plaque aggregation have provided important insights into the pathobiology of AD and there is consensus that they are representative models of the asymptomatic stage of AD (*4*), although it is accepted that they are incomplete models of the disease. Mice expressing mutant forms of human *APP* recapitulate many aspects of cerebral Aβ accumulation seen in the human disease, including associated neuroinflammation, synaptic dysfunction and vascular pathology (*5*). Characterization of the molecular events taking place during Aβ aggregate accumulation in the brain is key to identifying signaling pathways that might be altered during AD and to unveiling potential biomarkers and therapeutic opportunities. Previous efforts to characterize murine models of AD at the transcriptional level have highlighted the role of inflammatory pathways in AD pathogenesis (*6, 7*). Recently, key studies have addressed the role of neuroinflammation using single-cell sequencing strategies to identify specific microglial sub-populations associated with AD (*8, 9*). Given the accumulation of proteins in the plaques, quantitative proteomics might also be a fundamental approach to understand protein-related changes in AD pathogenesis. Unfortunately, to date, these studies have been limited to single time points and decoupled from gene-expression data (*10–14*). However, it is clear that combining transcriptional and proteomics data provides key insights into aging processes in rats (*15*).

Despite many advances in the characterization of different physiological motifs involved in AD onset and progression, our understanding of the events that trigger the disease are limited. Consequently, the only medical treatments currently available for AD are purely symptomatic and hardly effective (*16*), and most developments aiming at modifying the biology of the disease have failed (*17*). The recurrent failures have triggered a debate about the deficiencies in diagnostic strategies, the choice of therapeutic targets and the design of the clinical trials (*18*). On the one hand, it seems clear that modulation of a single target, even with a highly efficient drug, is unlikely to yield the desired outcome, and there is a growing perception that we should increase the level of complexity of our proposed therapies from a gene-centric to a systems view (*17*). Another aspect that has become apparent is that we have been studying AD and attempting to develop therapeutics against it at advanced stages of the disease when it is virtually impossible to revert the brain damage already caused (*19, 20*). We therefore need to focus on much earlier phases, ideally even before the first clinical symptoms appear or when cognitive impairment is still mild (*18*).

Unbiased genome-wide data-driven approaches may reveal new therapeutic opportunities, providing a global perspective beyond individual targets. For example, gene-expression data arising from a pathological phenotype can be seen as a disease-specific *signature* and used to find compounds able to restore a healthy state (*21*). Indeed, this approach has been successfully used to identify drugs targeting obesity (*22*), osteoporosis (*23*) and aging (*24*) at preclinical stages. Recently, we extended the idea of using small molecules to mimic or revert biological signatures beyond transcriptional profiles and demonstrated the capacity of our approach to identify compounds able to revert specific expression alterations of AD genes (e.g., BIN1, GRIND2D, etc.) in APP^V717F^ and PSEN1^M146V^ SH-SY5Y cells (*25*). However, despite its potential, this approach to revert global signatures has been validated only in cell cultures and thus its *in vivo* relevance is still uncertain.

Here we present a comprehensive characterization of three murine AD models at phenotypic and molecular levels, including transcriptomic and proteomic profiles. We explore the molecular changes associated with AD onset, progression and advanced stages, and correlate them with cognitive status and Aβ accumulation in order to study the dynamics of the disease and compare the changes observed with healthy aging processes. Moreover, to pinpoint potential instances of protein aggregation, we examine those cases in which transcript levels and protein abundance are decoupled. Finally, we use the derived molecular profiles to identify approved drugs able to revert the specific AD signatures and we study their effects at phenotypic and molecular levels *in vitro* and *in vivo*.

## Results

### Characterization of cognitive impairment in three AD mouse models

To gain an understanding of the dynamics of AD progression, we characterized different AD mouse models at the phenotypic and molecular level, at three representative stages of the disease (onset, progression and advanced). As primary models, we used two *App* mutated knock-in mouse versions widely used in AD preclinical research and developed to avoid potential artefacts of transgene overexpression (*26*). *App^NL-F^* and *App^NL-G-F^* mice contain a humanized Aβ sequence with the Swedish “NL” (KM670/671NL) and Beyreuther/Iberian “F” (I716F) mutations. In addition, the *App^NL-G-F^* model contains the Arctic mutation “G” (E693G) in the Aβ sequence (*27*). To complement these models, we also characterized the classical 3xTg-AD mice, which overexpress mutated human APP (*APP^KM670/671NL^*) and Tau (*MAPT^P301L^*) proteins in a *Psen1^M146V/M146V^* background (*28*).

To determine the disease stage in each AD model, we first evaluated the cognitive status of *App^NL-F^* and *App^NL-G-F^* mice of different ages using the novel object recognition (NOR) test, which is substantiated in the innate preference of mice for novelty (i.e., if the mouse recognizes a familiar object, it will spend most of its time exploring the novel object) (*29*). We used 3-, 9- and 18-month-old (mo.) *App^NL-F^* mice and 3-, 6- and 9-mo. *App^NL-G-F^* mice as representatives of different disease stages (*27*). We found that 18-mo. *App^NL-F^* mice spent a similar time exploring the novel object and familiar object (47.3 ± 11.1%), while their *App^wt^* counterparts spent 66.6 ± 7.5% of time exploring the novel object. Younger *App^NL-F^* mice (i.e., 3 and 9 mo.) did not show significant differences in object exploration time, thereby confirming that this AD model has memory defects only at advanced ages (Fig. 1a). In the case of the *App^NL-G-F^* mice, 6- and 9-mo. animals were not able to discriminate the novel object (53.2 ± 9.9% and 53.7 ± 9.9%, respectively), thereby indicating earlier cognitive impairment than the *App^NL-F^* mice (Fig. 1a). As expected, *App^wt^* animals spent more time on the novel object at all the ages tested. (63.6 ± 4.7%, 60.0 ± 12.2% and 61.4 ± 10.7% in 3-, 6- and 9-mo. mice, respectively).

**Figure 1.**
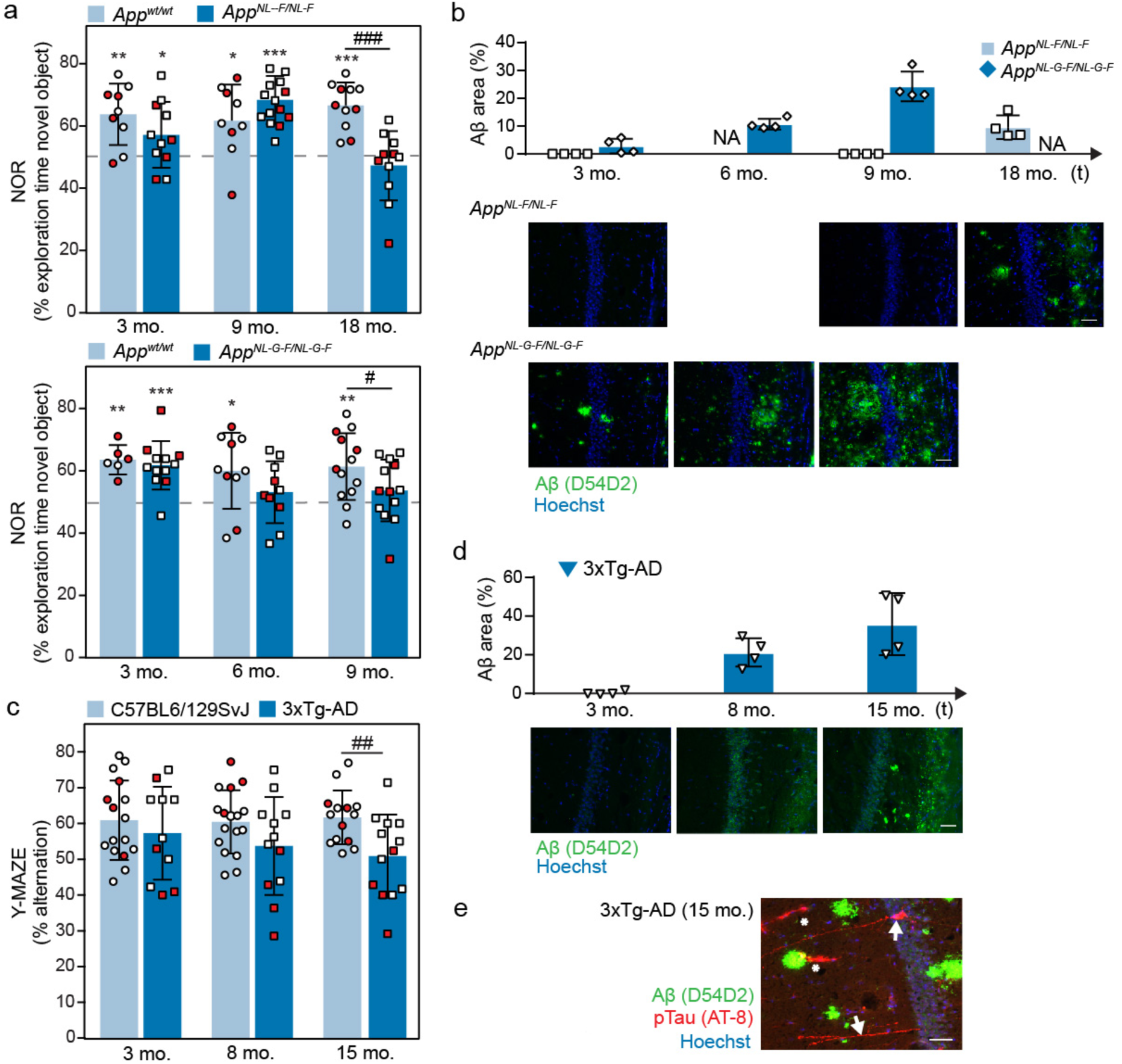
Behavioral and histological characterization of mouse AD models. **(a)** Top, novel object recognition (NOR) test performed on *App^wt^* (light blue; 3 mo. n= 9; 9 mo. n= 9; 18 mo. n= 11) and *App^NL-F^* (dark blue; 3 mo. n= 11; 9 mo. n= 15; 18 mo. n= 11) mice. Bottom, NOR test performed on *App^wt^* (light blue; 3 mo n= 6; 6 mo n= 10; 9 mo n= 13) and *App^NL-G-F^* (dark blue; 3 mo n= 12; 6 mo n= 10; 9 mo n= 13) mice. Mean±SD of the % of time exploring the novel object is shown. One-sample t-test versus a hypothetical value of 50 (* P < 0.05, ** P < 0.005, *** P < 0.0005) and unpaired Student’s t-test (# P<0.05; ### P < 0.0005) analysis are shown. Red dots indicate the results of the animals selected for histological and molecular profiling analyses. **(b)** Quantification and representative microphotographs of the CA1 region of the hippocampus of brain sections from 3-, 9- and 18-mo. *App^NL-F^* mice or from 3-, 6- and 9-mo. *App^NL-G-F^* mice stained with an anti-Aβ antibody (green) and Hoechst dye (blue) (n = 4 for each condition). Scale bars represent 50 µm. Photographs for both models were taken under identical conditions and the percentage of area with Aβ positive staining was quantified as indicated in the methods section. Mean±SD of *App^NL-F^* (light blue) and *App^NL-G-F^* (dark blue) mice are shown. NA: not available. **(c)** Y-maze test performed on mixed non-transgenic C57BL6/129SvJ mice (light blue; 3 mo. n= 16; 8 mo. n= 17; 15 mo. n=14) and 3xTg-AD transgenic mice (dark blue; 3 mo. n= 11; 8 mo. n= 12; 15 mo. n=13). Mean±SD of the % of alternation are shown. Unpaired Student’s t-test was performed (## P < 0.05). Red dots indicate the results of the animals selected for histological and molecular profiling analyses. **(d)** As in (a), sections from 3-, 8- and 15-mo. 3xTg-AD mice were stained with an anti-Aβ antibody (green) and Hoechst dye (blue) (n=4 for each condition). Scale bars represent 50 µm. Mean±SD are shown. **(e)** Representative staining of Aβ (green), phosphorylated Tau (red) and nuclei (blue) in a brain section of a 15-mo. 3xTg-AD mouse (n=4). White arrows indicate neurons with aggregated phosphorylated Tau. Asterisks indicate non-specific staining. Scale bar represents 50 µm.

We next selected samples of representative female mice (shown in red in Fig. 1a; see Methods for details) and quantified the Aβ aggregates in the hippocampal region of the two knock-in models (Fig. 1b). We observed that 18-mo. *App^NL-F^* mice presented small plaques with an Aβ-positive area of 9.6 ± 4.3%, whereas 3- and 9-mo. *App^NL-F^* mice did not show significant Aβ area staining. On the other hand, all 3-, 6- and 9-mo. *App^NL-G-F^* mice showed a progressive increase in Aβ staining (2.8 ± 2.6%, 10.7 ± 1.9% and 24.3 ± 5.3%, respectively). As expected, the corresponding *App^wt^* mice did not show Aβ staining (data not shown). Overall, these results were in line with the initial characterization of these mouse models (*27*), and comparison of hippocampus Aβ plaque accumulation with the results of the NOR test (Fig. 1a) pointed to the presence of a threshold of Aβ pathology from which these mice start to develop cognitive impairment (18 months for *App^NL-F^* mice and 6 months for *App^NL-G-F^* mice).

To complement the two knock-in models, we also characterized cognitive status (Fig. 1c) and Aβ plaque formation (Fig. 1d) in 3-, 8- and 15-mo. 3xTg-AD mice in a Y-maze spontaneous alternation experiment, which also evaluates hippocampal defects by measuring the willingness of mice to explore new environments. As expected, 8- and 15-mo. 3xTg-AD mice showed a progressively reduced percentage of alternation in the Y-maze (53.7 ± 13.7% and 50.9 ± 11.6%, respectively) compared with wild-type controls, which showed around 60% of alternation at all ages (Fig. 1c), suggesting cognitive impairment of the former. Although intracellular staining of Aβ was detected at 8 months, dense Aβ plaques were only clearly visible at 15 months of age (Fig. 1d). Note that the 3xTg-AD model has the unique characteristic of the concomitant manifestation of both plaques and tangles at late ages due to the overexpression of a mutated form of Tau (*MAPT*^P301L^). Indeed, we confirmed the presence of intracellular hyper-phosphorylated Tau in neurons of 15-mo. mice (Fig. 1e).

### Characterization of the gene expression and protein abundance associated with AD

To identify molecular changes associated with these pathological features, we performed a comprehensive parallel quantification of gene expression levels and protein abundance in the dissected hippocampi of the *App^NL-F^*, *App^NL-G-F^* and 3xTg-AD mice previously analyzed. More specifically, we used half the hippocampi for RNA sequencing of 3-, 9- and 18-mo. *App^NL-F^* mice; 3-, 6- and 9-mo. *App^NL-G-F^* mice; and 3-, 8- and 15-mo. 3xTg-AD mice (n=4 per genotype and time point), together with the corresponding *App^wt^* and C57BL6/129SvJ controls. Overall, we measured the expression levels of 21,950 protein-coding genes. Additionally, we used the other half of these hippocampi for proteomics studies, based on the 8-plex iTRAQ labeling method (*30*) and LC-MS/MS, we were able to quantify a total of 8,732 unique protein groups, corresponding to 6,837, 7,938 and 7,473 proteins from *App^NL-F^*, *App^NL-G-F^* and 3xTg-AD mice, respectively. A detailed description of the protocols is provided in the Methods section, and the complete datasets are available in GEO (*31*) GSE168431 and PRIDE (*32*) PXD024538.

To identify the changes in transcriptional and protein abundance associated with each disease stage, we first performed a differential abundance analysis comparing the AD genotypes at indicated ages with their wild-type counterparts, applying several False Discovery Rate (FDR) thresholds (Fig. 2a). We also compared different ages within the same genotype to evaluate changes associated with disease progression for all three models (Fig. 2b). Consistent with the phenotypes described in Fig.1, 6- and 9-mo. *App^NL-G-F^* mice showed the largest effects on gene/protein down- and, specially, up-regulation (150 genes and 51 proteins for 6-mo. mice, and 295 genes and 64 proteins for 9-mo. mice; n=4, FDR<5% and |logFC|>0.5 and 0.25). Interestingly, few changes were found in the 6- vs. 9-mo. comparison (Fig. 2b; 18 genes and 0 proteins; n=4, FDR <5% and |logFC|>0.5 and 0.25). This observation thus indicates that most of the changes take place between 3 and 6 months of age, when Aβ plaques became more evident and cognitive impairment appeared (Fig. 1a and 1b). On the other hand, despite the small Aβ plaques observed in 3-mo. *App^NL-G-F^* mice (Fig. 1b), no detectable changes in transcript or protein abundance were observed with respect to their wild-type counterparts (Fig. 2a). Similarly, 18-mo. *App^NL-F^* mice did not show differences with their age-matched wild-type controls (Fig. 2a). However, an age comparison (Fig. 2b; 3- vs. 18-mo. *App^NL-F^*) identified age-dependent differentially expressed genes/proteins (107 and 72, respectively; n=4, FDR <5% and |logFC|>0.5 and 0.25). When analyzing the genotype-dependent changes in the 3xTg-AD model, we already observed significant gene expression and protein abundance differences at 3 months of age (Fig. 2a), even before the first signs of Aβ pathology appeared in the hippocampus of these mice (Fig. 1d). Moreover, in contrast to the *App^NL-G-F^* model, the number of differentially expressed genes/proteins did not increase with disease progression (Fig. 2a). Indeed, a principal variance component analysis (PVCA) showed that the genotype accounted for 48% and 19% of the variance observed in the transcriptomics and proteomics data for this model, compared to the 17% and 13% observed in *App^NL-G-F^* and the 8% and 4% observed in *App^NL-F^* models (Fig. S1b). We attribute this effect to the wild-type controls used with the 3xTg-AD model, since one of the drawbacks of the 3xTg-AD model is that it has to be kept in a mixed background, as backcrossing may affect the initially reported phenotype (*33*). Therefore, similar mixed background mice (as recommended by The Jackson Laboratory) were used as controls. This effect does not influence the comparisons across ages in the 3xTg-AD model, where we observed, as expected, an increase in dysregulated genes/proteins as AD advanced (Fig. 2b).

**Figure 2.**
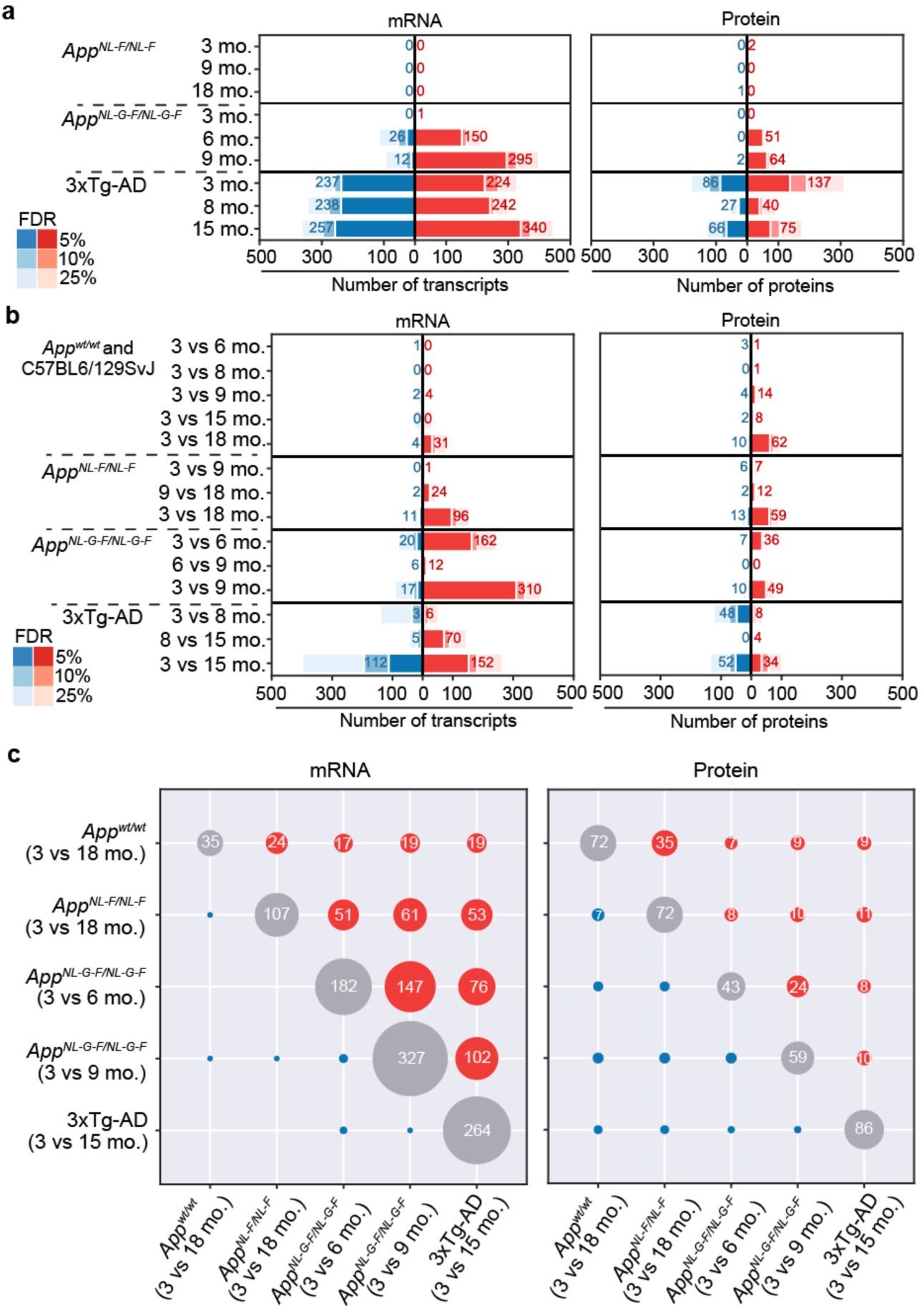
Differential molecular profiling and AD progression. **(a)** Number of differentially expressed genes (left panel) and proteins (right panel) comparing the three mouse AD models (*App^NL-F^, App^NL-G-F^* and 3xTg-AD) at different ages with their corresponding, age-matched, wild-type controls. Red and Blue represent up and downregulated genes/proteins, respectively. Bars indicate the number of significant genes/proteins that changed between conditions at different thresholds of False Discovery Rate (FDR) and with an absolute Log Fold Change (LogFC) larger than 0.5 for mRNA and 0.25 for proteins. Numbers depicted in the figure correspond to FDR<5%. N=4 for all the conditions. **(b)** Number of differentially expressed genes (left panel) and proteins (right panel) comparing each genotype at the different ages. N=4 for all the conditions. **(c)** Overlap between differentially expressed genes (left panel; absolute LogFC>0.5, FDR<5%) or proteins (right panel; absolute LogFC>0.25, FDR < 5%) comparing AD mouse models and disease stages.

Interestingly, we observed a significant degree of overlap between the upregulated genes and proteins in the three different models, thereby suggesting that, despite the notable differences between models, there are common molecular patterns across them (Fig. 2c). For instance, the strongest overlap across models occurred between *App^NL-G-F^* (3- vs. 9-mo.) and 3xTg-AD (3- vs. 15-mo.) mice, which shared 102 upregulated genes (odds ratio (OR) 247.53, Fisher’s exact test p-value 7.58·10^-164^) and 10 proteins (OR 62.47, p-value 6.45·10^-14^). On the other hand, *App^NL-F^* (3- vs. 18-mo.) also shared a significant number of upregulated genes and proteins with both *App^NL-G-F^* (3- vs. 9-mo.), with 61 genes and 10 proteins (OR 172.13 and 29.22; p-value 1.37·10^-94^ and 3.96·10^-11^) and 3xTg-AD (3- vs. 18-mo.), with 53 genes and 11 proteins (OR 316.27 and 53.82; p-value 1.12·10^-95^ and 1.99·10^-14^). As for the downregulated genes and proteins, the changes were much less evident and specific for each model. Indeed, only 17 genes and 10 proteins were significantly downregulated in *App^NL-G-F^* mice (3- vs. 9-mo.), and the only significant overlap between downregulated genes and proteins corresponded to the *App^NL-F^* (3- vs. 18-mo.) and *App^wt^* (3- vs. 18-mo.) mice, which involved 7 proteins (OR 2250.89 and p-value 4.68·10^-18^).

Additionally, analysis of the *App^wt^* mice corresponding to the knock-in models allowed us to study the relationship between healthy aging and AD progression at the molecular level. Comparison of “old” (18-mo.) with “young” (3-mo.) wild-type mice revealed the upregulation of 31 genes and 62 proteins and the downregulation of 4 genes and 10 proteins (Fig. 2b; Table S1; FDR<5% and |logFC|>0.5 and 0.25). At the transcriptional level, we detected the upregulation of known microglial markers such as *Cst7*, *Clec7a* or *Lyz2* (Fig. S2a), thereby suggesting an increase in the inflammatory component in the aged hippocampus. This finding is in line with the recent characterization of gene expression in aged microglia at single-cell level (*8, 34, 35*), and also with the general accepted role of inflammation as one of the hallmarks of aging (*36*). Of note, a significant number of these genes were also upregulated in other comparisons involving younger AD mice (e.g. 17 of the 21 genes were already upregulated in 3- vs. 6-mo. *App^NL-G-F^* animals, OR 215.73 and p-value 3.48·10^-30^; Fig. 2c and Fig. S2a), thereby suggesting that Aβ pathology promotes features of molecular aging. However, the observation that 18-mo. *App^wt^* mice did not present cognitive deficits (Fig. 1b) confirms that AD is not merely accelerated aging, and some differences must exist with *App^NL-F^* mice, which indeed showed cognitive impairment at 18 months of age (Fig. 1b) despite the lack of significantly altered genes or proteins (Fig. 2a). Direct comparison between *App^wt^* (3- vs. 18-mo.) and *App^NL-F^* (3- vs. 18- mo.) mice indicated a similar trend of up- and down-regulated genes (Fig. S2b), although we identified a set of genes with accentuated changes in *App^NL-F^* mice (Table S2). Among these, we found a number of chemokines (*Ccl6*, *Ccl3* or *Ccl5*) and also markers of microglial activation (*Cd14*, *Tyrobp*). Interestingly, *Ccl3* has been found to impair mouse hippocampal synaptic transmission, plasticity and memory, and polymorphisms in this gene increase the risk of AD in humans (*37*).

While there was a significant overlap between the transcriptional changes in 3xTg-AD mice (3- vs. 15-mo.) and those observed in the *App^NL-G-F^* model (76 with 3- vs. 6-mo. comparison; 102 with 3- vs. 9-mo. comparison; Fig. 2c), we would expect that some of the changes observed in the 3xTg-AD model could be attributed to the presence of Tau fibrils (Fig. 1e). Therefore, we directly compared changes in 3xTg-AD mice (3- vs. 15-mo.) with *App^NL-G-F^* mice (3- vs. 9- mo.) to identify 3xTg-AD-specific changes (Fig. S2c; Table S3). The 3xTg-AD model showed an accentuated upregulation of *Klk6* and *Lcn2*, which have been evaluated as possible markers of AD and vascular dementia, respectively (*38, 39*). We also observed an increased upregulation of Serpina3n protein, a serine protease inhibitor previously identified in human amyloid deposits (*40*) and dysregulated in prion diseases (*41*), which might indicate a role for Tau aggregates in the metabolism of these diseases. Additionally, we found the upregulation of *Alox12b*, which encodes a lypoxigenase enzyme, only in the 3xTg-AD model. This observation is consistent with recent findings in a Tau mouse model (*42*) and the potential role that lipoxygenases might have in Tau metabolism (*43*). Of note, we consistently detected the downregulation of extracellular matrix genes (*Col1a2*, *Col3a1*) and proteins (Col6a1), as well as proteins linked to adhesion or actin function, such as Flna, Palld or Fbln5 (Fig. 2c; Table S3). These observations thus support the hypothesis that hyper-phosphorylated Tau compromises the integrity and function of the blood–brain barrier in the 3xTg-AD mouse model (*44*).

### Physiological aging and AD progression

To gain further insight into the common molecular events that take place in response to pathogenic *App* mutations, we integrated the two *App* knock-in mouse models (*App^NL-F^* and *App^NL-G-F^*) and their respective *App^wt^* mice to analyze the entire experiment as a whole. To this end, and in order to capture the temporal dynamics of the transcriptional changes associated with physiological aging and AD progression, we considered the age of the animals as a continuous variable. We measured the linear association of gene expression with age for each genotype and identified a series of up- and down-regulated genes in *App^NL-G-F^* mice with respect to *App^wt^* that we named *AD signatures* (*AD-UP* and *AD-DW*, respectively; Table S4). Finally, we compared the relative expression of AD signature genes across all the mouse models (Fig. 3a and Fig. 3c, left panels) and observed that a number of up- and down-regulated genes followed similar trends in the comparisons of 3- vs. 15-mo. and 8- vs. 15-mo. 3xTg-AD mice (Fig. 3a and 3c, left panels). Indeed, the *AD-signatures* derived from the *App* knock-in mice were significantly enriched in the 3xTg-AD transcriptional changes (Fig. S3a), indicating that these signatures are common to the three AD models. Moreover, we found that the derived AD signatures included 28 genes defined as AD risk factors in human genetic studies (*45*) (Odds ratio (OR) 3.16 and Fisher’s exact test *p*-val 6.29·10^-7^), 23 of them in the *AD-UP signature* and 5 in the *AD-DW signature* (Fig. S3b and Table S4).

**Figure 3.**
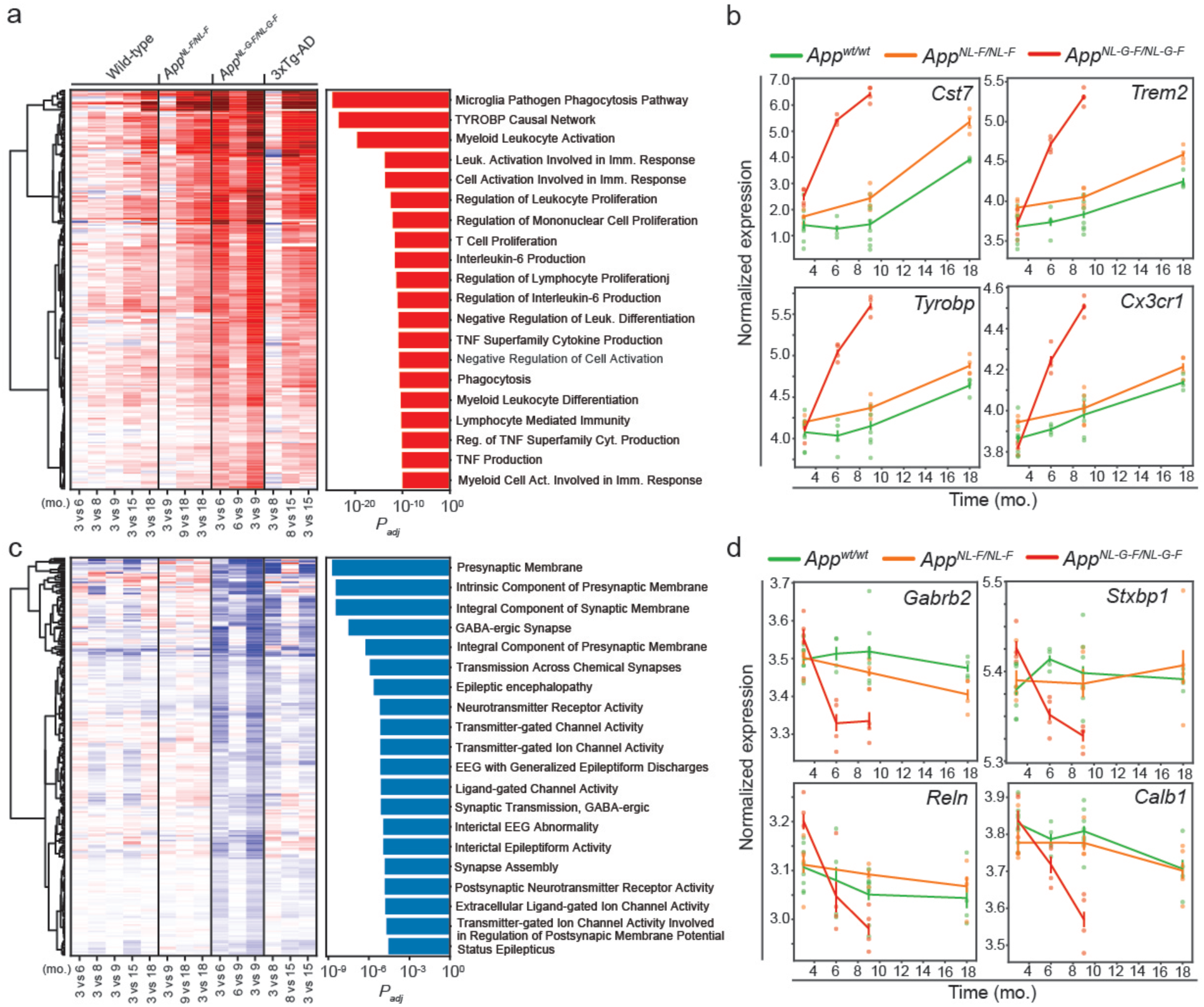
Identification and functional annotation of AD signatures. **(a)** Heatmap showing the progression of the transcriptional changes in the top-250 genes showing a positive genotype by age interaction (*AD-UP signature*). Each gene is represented by a row of colored tiles, the color representing the RNA expression level for the indicated condition by column (red, upregulated; blue, downregulated). The bar plot on the right indicates the top-20 significant pathways corresponding to the functional enrichment analysis of these top-250 genes, indicating the adjusted p-value on the X-axis. **(b)** Normalized gene expression at different time points expressed in months (mo.) for four examples of the *AD-UP signature* (*Cst7*, *Trem2*, *Tyrobp* and *Cx3cr1*) in the *App^NL-G-F^* (red line), *App^NL-F^* (orange line) and *App^wt^* (green line) mice. N=4. **(c)** As in (a), progression of the transcriptional changes in the top-250 genes showing a negative genotype by age interaction (*AD-DW signature*) and functional enrichment analysis of these genes. **(d)** As in (b), four examples (*Gabrb2*, *Stxbp1*, *Reln* and *Calb1*) of the *AD-DW signature* are shown. N=4.

We performed a functional enrichment analysis of AD signatures and found that the top-20 upregulated pathways corresponded exclusively to the activation of the immune system, including phagocytosis or cytokine production pathways (Fig. 3a, right panel). Indeed, many of the top upregulated transcripts were directly associated with microglia activation (Fig. 3b), including *Tyrobp* and its putative receptor *Trem2*, whose variants are associated with an extraordinarily increased risk for AD in humans (*46*). Our AD signatures also included 46 disease-associated microglia genes (46 of 382; OR 23.33, p-val 4.39·10^-37^), which have been associated with microglial changes linked to pathological insults (*8*), and 38 plaque-induced genes (38 of 56; OR 255.62, p-val 2.4·10^-64^) previously identified in a spatial transcriptomics characterization of the cellular response to amyloidosis (*47*). These findings are therefore consistent with those of other studies using distinct approaches.

One of the most upregulated genes in this *AD-UP signature* was *GFAP*, which was also upregulated at the protein level (Table S4), and we confirmed by immunofluorescence in the *App^NL-G-F^* model that it activates astrocytes around Aβ aggregates (Fig. S4a). GFAP is an astrocyte marker, and it has been reported to accumulate in the brains of AD patients (*48*). Additionally, six genes in the *AD-UP signature* (e.g. *Cd14*, *Gfap*, *Aspg*, *S1pr3*, *Ggta1* and *Serpina3n*) were also found to be involved in astrocyte activation (*49*), and we found others (e.g., *Vim*) significantly upregulated in both *App^NL-G-F^* and 3xTg-AD mice (Fig. S4b).

Overall, our *AD-UP signature* was able to recapitulate known activation pathways (microglia and astrocyte), including a significant number of known AD genetic risk factors, in response to pathological accumulation of amyloid plaques, in agreement with previous studies. Immune response to amyloid accumulation is an important part of the pathology of the disease, but the convenience or the timing for blocking or stimulating this response as a therapeutic option is still arguable (*50*).

On the other hand, functional enrichment analysis of the *AD-DW signature* highlighted defects on synapse transmission, synapse membrane components, neurotransmitters or the GABA-ergic signaling system (Fig. 3c, right panel). For example, several GABA-ergic receptors such as *Gabra1*, *Gabra3* or *Gabrb2,* as well as neuroprotective *Calb1* or *Reln* genes, were downregulated (Table S4 and Fig. 3d). As these molecular defects may reflect the cognitive impairment observed in the *App^NL-G-F^* mice at 6 and 9 months of age, reverting some of these molecular changes could offer therapeutic opportunities (*51*).

### Identification of proteins associated with Aβ plaques

Access to both transcriptomics and proteomics data allowed us to analyze the degree of correlation between the two datasets for proteins that were significantly up- or down-regulated in the different age and AD-model comparisons (Fig. 4a and Fig. S5a). Of note, the changes observed at the protein level were strongly correlated with transcriptional changes in the AD models (*App^NL-G-F^*, *App^NL-F^* and 3xTg-AD), indicating that many of the proteomic changes happening during Aβ pathology are triggered and regulated at the transcriptional level. This correlation was much weaker in physiological aging (3- vs. 18-mo. *App^wt^*), where we observed a marked accumulation of proteins that was uncoupled from transcriptional changes. Interestingly, the proteins that accumulated in aged mice (3- vs. 18-mo. *App^wt^* and *App^NL-F^* and 3- vs. 15- mo. 3xTg-AD) tended to have longer lifespans compared to those that accumulated in *App^NL-G-F^* mice, thereby suggesting that different mechanisms regulate protein homeostasis depending on the age (Fig 4a, Fig S5). We then analyzed, from a functional perspective, the proteins whose higher levels were not explained by changes at the mRNA level. We found common pathways related to pH regulation and glycogen metabolism in most aged models (i.e. 3- vs. 18-mo). *App^wt^* and *App^NL-F^* mice, while proteins were involved in Aβ formation or metabolic processes in the most aggressive models (Fig. 4b), thereby suggesting that some of the proteins interacting with Aβ, or its precursor App may be accumulating. For instance, we found that the protein levels of Itm2c were increased in the *App^NL-G-F^* mice while its transcripts remained stable (Fig. 4c). Indeed, this protein has been found to co-localize with Aβ plaques in mouse and human brain samples (*52, 53*). We also detected an acute increase in Ifit3 protein levels, compared to mRNA levels, in 6-mo. *App^NL-G-F^* mice (Fig. 4c). In a previous study, we proposed that Ifit3 might physically interact with App (*54*), making its aggregation in amyloid plaques plausible. IFIT family members regulate immune responses and restrict viral infections through a variety of mechanisms, including the restriction of RNA translation or binding to viral proteins (*55*). Therefore, we used immunofluorescence to stain Ifit3. We observed that it formed plaque-like structures that co-localized with Aβ plaques in the brains of 6-mo. *App^NL-G-F^* mice but these structures were absent in *App^wt^* control mice (Fig. S5). Interestingly, we also identified a sharp increase in Synaptotagmin-11 (Syt11) protein levels in the *App^NL-G-F^* model, while its transcripts decreased (Fig. 4d). Syt11 is genetically linked to risk of Parkinson’s disease (*56*) and it is a substrate of PRKN (encoded by *PARK*2), an E3 ubiquitin ligase that is often mutated in familial cases of Parkinson’s disease (*57, 58*). In mice, Syt11 deficiency in excitatory glutamatergic neurons impairs synaptic plasticity and memory (*59*), thereby suggesting that Syt11 dysregulation contributes to AD-associated cognitive decline. Analysis of the brain sections of 3-, 6- and 9-mo. *App^wt^* and *App^NL-G-F^* mice revealed Syt11 dense stains accumulating in the latter, especially at 6 and 9 months of age (Fig. 4e). Importantly, we found that Syt11 co-localized with Aβ plaques in *App^NL-G-F^* mice (Fig. 4f), thereby suggesting that accumulation of Syt11 in amyloid plaques contributes to the AD pathology. Moreover, a proximity-ligation assay (PLA) showed that endogenous Syt11 and App proteins interacted in human neuron-like SH-SY5Y cells (Fig. S6). Although we did not observe Syt11 dense stains in the brains of *App^NL-F^* or 3xTg-AD mice at the ages tested (18 and 15 months of age, respectively), a recent proteomics analysis in the cortex of the FADx5 AD mouse model also found Ifit3 and Syt11 among the most upregulated proteins at advanced ages (*60*). This finding indicates that our results could be extrapolated to other models.

**Figure 4.**
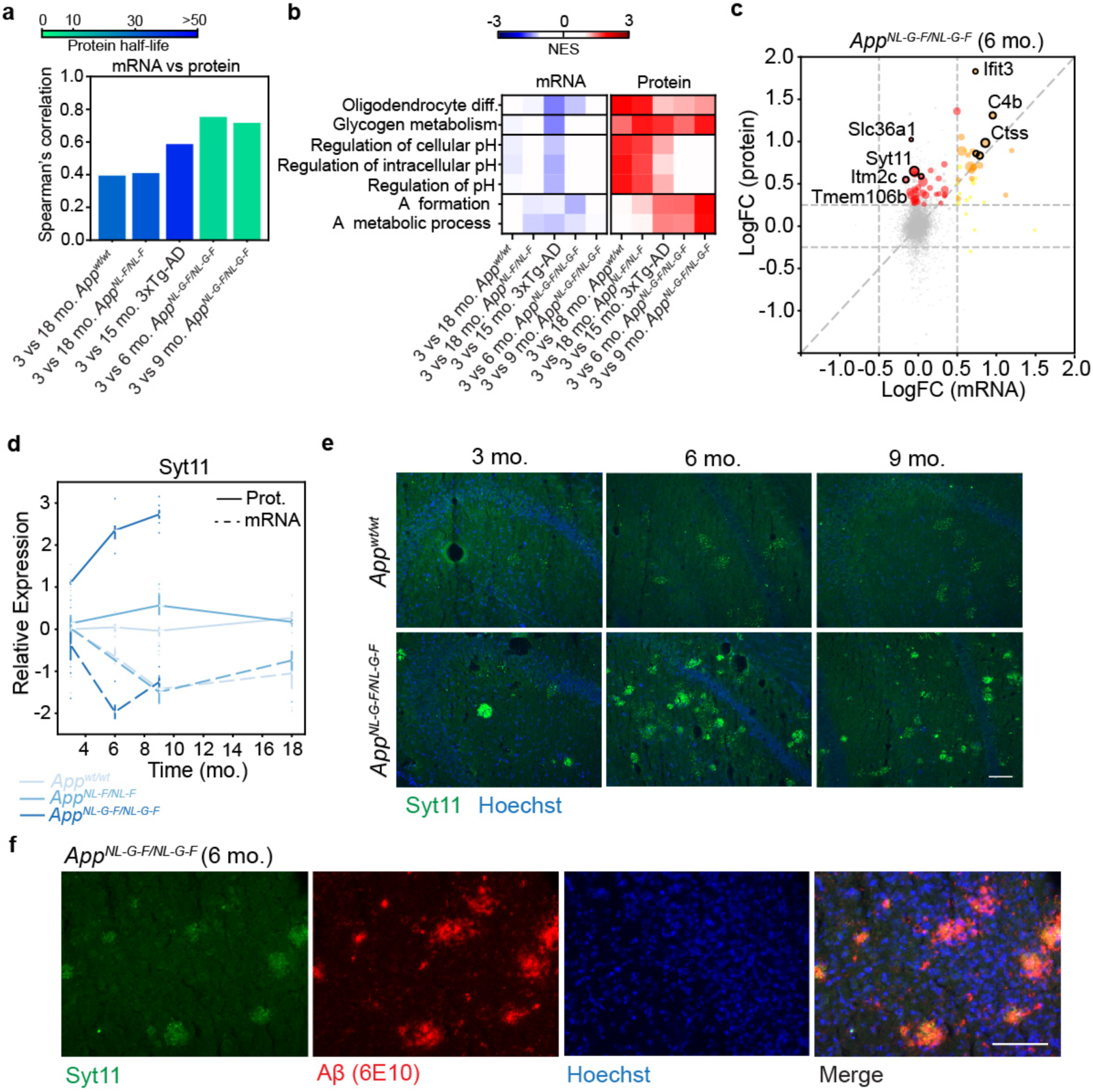
Identification of Aβ-aggregated proteins through comparison of mRNA and protein levels. **(a)** Spearman’s rank correlation of significantly altered proteins with respect to their mRNA expression in the indicated comparisons. Bar colors indicate the mean half-life of the proteins as defined by (*96*). **(b)** Discordant upregulation of GO biological processes and Reactome pathways at protein level that is not explained by the upregulation of the same genes at mRNA level. **(c)** Scatter plot depicting the logarithm of the Fold-Change (LogFC) of mRNA (X-axis) and protein (Y-axis) for the *App^NL-G-F^ vs. App^wt^* comparison at 6 mo. Proteins whose LogFC(protein)>0.25 and their corresponding LogFC(mRNA)>0.5 are shown in orange while those with a LogFC(protein)>0.25 and LogFC(mRNA)<0.5 are shown in red. Dot size is proportional to the negative logarithm of the adjusted p-value. **(d)** Syt11 protein (continuous line) and mRNA (dashed line) levels at different time points relative to the *App^wt^* at 3 mo are shown for the *App^NL-G-F^* (strong blue), *App^NL-F^* (medium blue) and *App^wt^* (corresponding to the *App^NL-F^* model; light blue) mice. N=4. **(e)** Representative microphotographs of the hippocampus of brain sections from 3-, 6- and 9-mo. *App^wt^* (top row) or *App^NL-G-F^* (bottom row) mice stained with an anti-Syt11 antibody (green) and Hoechst dye (blue) (n = 3 for each condition). Scale bars represent 100 µm. **(f)** Representative micrographs of a brain section of a 6-mo. *App^NL-G-F^* mouse stained with an anti-Syt11 antibody (green), the anti-Aβ antibody 6E10 (red) and Hoechst dye (blue). Scale bars represent 100 µm (n=3).

### Identification of approved drugs with the potential to revert AD signatures

Next, we sought to identify small molecules with the capacity to “revert” the transcriptional traits of AD mice, potentially ameliorating the disease phenotype. As a chemical space, we considered the over 800,000 bioactive compounds included in the Chemical Checker (CC) (*25*). The CC provides different types of bioactivity descriptors (a.k.a. *signatures*) for each compound, including transcriptional and target profiles gathered from the major chemogenomics databases, as well as the more conventional descriptors of chemical structure. We can then use these signatures to *connect* small molecules to desired outcomes observed in phenotypic experiments such as genetic perturbation assays, as popularized in the context of transcriptomics by the Connectivity Map initiative (*61*). Indeed, we proved that CC signatures could be used to identify compounds able to revert the transcriptional changes induced by AD mutations (PSEN1^M146V^ and APP^V717F^) in SH-SY5Y cells (*25*). We now explored whether the capacity to prioritize compounds that revert molecular signatures could be translated to *in vivo* models, in which the phenotypic effects of this reversion can be measured.

We devised five distinct strategies to query the CC compound signatures based on known AD pharmacology (queries 1-3) and on the transcriptional profiles observed in our AD mouse models (queries 4-5). More specifically, we looked for compounds chemically similar to drugs clinically tested for AD (*62*) or with similar mechanisms of action (1), or compounds that elicited similar transcriptional responses to these drugs (2). In addition, we searched for molecules that could bind putative AD targets, as defined by OpenTargets (*63*) (3). Moreover, we used the CC gene expression signatures to identify compounds that trigger transcriptional changes in the opposite direction to those observed in our AD mice models (4). Likewise, we looked for shRNA experiments in the LINCS L1000 to identify knock-downs that could also revert our AD signatures, and we used them as putative AD targets to recall further candidate compounds from the CC (5). Finally, we computed blood-brain barrier penetration (BBBP) and Aβ affection scores for all the compounds, based on machine-learning models. More details on these search procedures are available in the *Methods* section. Overall, by using these five queries we shortlisted 1% of the CC, yielding a collection of 8,250 candidates (Table S5). Of these, and in order to avoid issues related to compound stability and solubility *in vivo*, we kept only those that are commercially available and those previously tested in mice.

Among the 8,250 candidate compounds, we found 125 highly ranked non-steroidal anti-inflammatory drugs (NSAIDs) targeting PTGS1 or PTGS2. NSAIDs are one of the most widely used types of medication, being prescribed as anti-inflammatories, analgesics and antipyretics. Although clinical trials have failed to demonstrate beneficial effects of NSAIDs in AD (*64*), recent articles suggest that some exert previously undescribed mechanisms of action that ameliorate AD pathology in mouse models (*65, 66*), thus supporting epidemiological studies reporting a potential protective effect of NSAIDs against AD (*67, 68*). Thus, we selected three NSAIDs (namely etodolac, fenoprofen and dexketoprofen) considering the scores obtained in all the described CC, and whose activity in AD had not been tested previously. Etodolac and fenoprofen are chemically similar to other NSAIDs clinically tested against AD (query 1) and they performed, in general, relatively well in all queries. Additionally, fenoprofen and dexketoprofen showed an interesting targets profile (query 3), and the former also presented a high potential for reverting the transcriptional changes observed in our AD models (query 4). All three compounds presented reasonable BBBP scores (>0.6).

The list also contained a number of approved anti-hypertensive agents, including structurally similar non-selective beta-adrenergic antagonists such as penbutolol, levobunolol, nadolol and bupranolol. Among these, penbutolol showed a strong aggregated score for targets (query 3) and good potential for reverting transcriptional signatures (query 4) from published profiles (*69*) and our models. We also selected other anti-hypertensive agents with different mechanisms of action, including bendroflumethiazide, which targets sodium reabsorption, and pargyline, a MAO-A inhibitor with a relatively high score in query 3 (AD targets profile). Out of the six compounds, bendroflumethiazide showed the best reversion score for our AD signatures, ranking 31st out of the 8,250 compounds sorted in the first instance, and its effects were also phenotypically similar to compounds tested in the clinics for anti-AD activity (query 2). We also calculated a reversion score for human AD-associated signatures, extracted from ((*69*); see Methods). Again, bendroflumethiazide showed an extremely high score (3.82), being the 37th best score among the 8,250 preselected compounds.

We assessed the effectiveness of the selected compounds on the *App^NL-G-F^* model, since it shows cognitive deficits at early stages. We treated 5-mo. mice with the different drugs for four weeks and, during the characterization of the models, we evaluated the effects of the drugs in the NOR test. We then collected hippocampi samples for further molecular analyses (Fig. 5a). As we had already observed, *App^NL-G-F^* mice treated with vehicle performed worse (48.6 ± 12.5% for vehicle 1 and 53.5 ± 15.7% for vehicle 2) than age-matched *App^wt^* mice in the NOR test (60.1 ± 10.7% for vehicle 1 and 58.0 ± 10.5% for vehicle 2) (Fig. 5b). On average, *App^NL-G-F^* mice treated with bendroflumethiazide (62.4 ± 18.4%), dexketoprofen (55.7 ± 6.1%), etodolac (57.5 ± 23.3%) or penbutolol (68.1 ± 15.5%) performed better than the corresponding control mice treated with vehicle. However, we did not appreciate any cognitive improvement in the mice treated with fenoprofen (47.3 ± 17.0%) or pargyline (49.6 ± 4.7%). Overall, four out of the six treatments assayed yielded NOR results comparable to those of wild-type animals, thereby pointing to the potential rescuing of the cognition impairment associated with *App* mutations.

**Figure 5.**
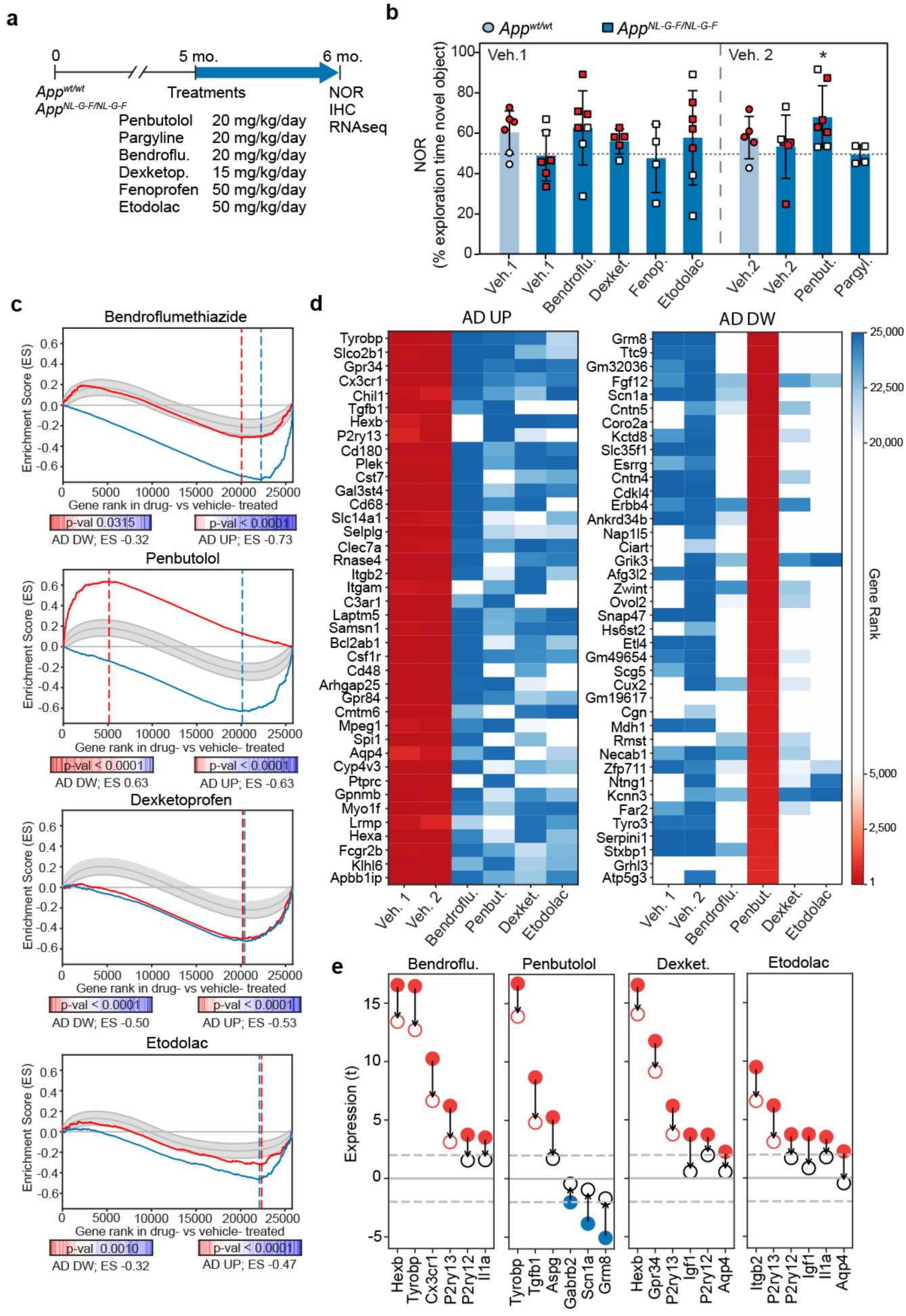
*In vivo* reversion of AD signatures. **(a)** Scheme of drug treatments and evaluation Novel object recognition (NOR) test of 6-month-old animals. *App^wt^* (circles; light blue) and *App^NL-F^* (squares; dark blue) mice were treated for 4 weeks with the indicated drugs. Mean±SD of the % of time exploring the novel object is shown (n= 4-7). One-sample t-test vs. a hypothetical value of 50 (* P < 0.05). Red points indicate the animals selected for RNAseq analysis. **(c)** Signature reversion. In the X-axis, genes are ranked by their differential expression in the comparison of drug- vs. vehicle-treated *App^NL-G-F^* mice. The Y-axis represents the running Enrichment Score (ES) performed for the *AD signatures* (*AD-UP*, blue; *AD-DW*, red) that were reverted upon treatment. Dashed vertical lines indicate the point at which the ES reaches its maximum deviation from zero, defining the leading-edge subset of genes that contribute the most to the enrichment result. The observed ES is compared to a null distribution of 10,000 randomized signatures of the same size (95% CI shown as a shaded gray area). For example, *AD-DW signature* genes (red line) tend to be ranked among the 5,000 most upregulated genes in response to penbutolol treatment (second panel), with an ES of 0.63, which is significantly higher than that of random signatures of the same size (p-value <0.0001). This is interpreted as a significant signature reversion. RNAseq data were obtained from n= 4 mice per condition. **(d)** Heatmaps showing the *reversion rank* of the top-40 genes belonging to the leading-edge of the *AD-UP signature* (blue; left panel) and *AD-DW signature* (red; right panel) in two or more treatments, sorted by the average row value. Data obtained from n=4 mice per condition. **(e)** Leading-edge genes of the *AD-UP* and *AD-DW signatures* reverted by the different treatments. We show a few genes in the AD signatures that are up- (red) or down- (blue) regulated (t-score) in the vehicle-treated *App^NL-G-F^* mice compared with vehicle-treated *App^wt^* animals (bold dots) or in the drug-treated *App^NL-G-F^* mice compared with vehicle-treated *App^wt^* animals (empty dots). RNAseq data were obtained from n=4 mice per condition.

In view of these results, we analyzed the gene expression changes in the hippocampi of *App^NL- G-F^* mice treated with the compounds that showed a beneficial effect in the NOR test (bendroflumethiazide, dexketoprofen, etodolac and penbutolol) and the corresponding controls (vehicle-treated *App^NL-G-F^* and *App^wt^* mice; Table S6). As expected, analysis of *App^wt^* and *App^NL- G-F^* controls showed similar transcriptional signatures compared with our previous characterization of the model (data not shown). Indeed, the four compounds significantly downregulated the expression of genes upregulated in the AD signature, thus being able to revert these characteristic transcriptional changes related to Aβ pathology. Of the four compounds, bendroflumethiazide and penbutolol showed the strongest effects (blue lines, Fig. 5c). Functional enrichment analysis of these reverted genes showed significant association with phagocytosis and activation of the immune response pathways (Fig. S7a), which is coherent with the strong component of immune system activation found in the signatures of the AD mouse models and the fact that these signatures were used to prioritize the selection of potential drugs. Genes such as *Cx3cr1* or *Tyrobp*, previously identified as part of the *AD-UP signature,* were partially downregulated in bendroflumethiazide-treated *App^NL-G-F^* mice compared with vehicle-treated mice (Fig. 5d and Fig. 5e), suggesting that this drug may inhibit microglia activation. Bendroflumethiazide is an anti-hypertensive drug that exerts its effect by inhibiting sodium reabsorption at the beginning of the distal convoluted tubule. Although Na^+^ influx has been linked to activation of the inflammasome (*70*), we could not find previous reports detecting or suggesting a potential anti-inflammatory role.

Although all four treatments reverted the tendency of upregulated genes in AD signatures, only penbutolol treatment also induced the reversion of a significant proportion of the *AD-DW signature* in *App^NL-G-F^* mice (Fig. 5c; red lines). Penbutolol prevented the Aβ-associated downregulation of genes such as the gamma-aminobutyric acid (GABA) receptor *Gabrb2* or the glutamate receptor *Grm8*, keeping their expression at physiological levels (Fig. 5d and Fig. 5e).

Next, we used immunofluorescence to quantify Aβ accumulation in the brains of treated *App^NL-G-F^* mice. The percentage of Aβ-positive area in the DG-CA1 hippocampal regions of *App^NL-G-F^* mice treated with penbutolol (5.1 ± 1.6%) was reduced compared with mice treated with the corresponding vehicle (6.3 ± 1.8%; Fig. S8a). As expected, *App^wt^* animals did not show Aβ staining. Other treatments did not seem to have an effect on Aβ accumulation when compared to the corresponding vehicles (data not shown). To further validate the effect of penbutolol, we treated neuron-like SH-SY5Y cells, which are known to recapitulate phenotypes related to neurodegenerative disorders (*71*). Indeed, we observed that penbutolol inhibited the secretion of Aβ_40_ (47.5 ± 15.5% at 25 µM) in a dose-dependent manner, while a minor effect was observed for Aβ_42_ secretion (78.8 ± 14.2% at 25 µM), without any observed toxicity (Fig. S8b). Additionally, we tested the compounds in genetically modified SH-SY5Y cells harboring the familial AD mutation *PSEN1^M146V/M146V^* (*25*), which increases Aβ_40_ (1.6 ± 0.3 fold) and Aβ_42_ (3.4 ± 0.2 fold) secretion (Fig. S8b). Like in wild-type cells, penbutolol inhibited the secretion of Aβ_40_ (67.2 ± 6.3% of DMSO control at 25 µM) in *PSEN1^M146V/M146V^* cells (Fig. S8b). Finally, we tested the effect of these compounds in the 7PA2 cell line, a well-established model for screening compounds targeting Aβ production (*72*). In accordance with the *in vivo* observation, penbutolol inhibited Aβ_40_ (16.1 ± 10.0% at 25 µM) and Aβ_42_ (30.9 ± 24.0% at 25 µM) at relatively low concentrations, showing a dose-dependent response. Fenoprofen showed a similar trend, although the inhibition of Aβ_40_ was milder (53.3 ± 4.7% at 500 µM), while the remaining four compounds did not exhibit any significant effect (Fig. S8c).

We had previously identified astrocytosis as an important component of the AD-associated molecular changes in our mouse models (Fig. S4) and, interestingly, *Gfap* was one of the genes whose expression was partially downregulated in response to penbutolol (Fig. S8d). Moreover, penbutolol was able to significantly revert the previously defined astrocytic signature (Fig. S8d). Also, the percentage of GFAP immunofluorescence positive area was slightly reduced in the brain sections of *App^NL-G-F^* mice treated with penbutolol (12.3 ± 3.7%) when compared with vehicle-treated mice (14.3 ± 4.3%; Fig. S8e) and, as expected, in both cases we observed more GFAP staining than in the corresponding age-matched *App^wt^* mice (8.8 ± 2.4% for vehicle 2).

Overall, our results consistently suggest that penbutolol inhibits Aβ production, accompanied by a reversion of the AD signature, both ameliorating inflammation, astrocytosis and loss of synaptic genes. Although the phenotype reversion is mild, optimized regime treatments (e.g. longer treatment, earlier treatment or improved delivery) might improve the results.

## Concluding Remarks

In this manuscript, we present the complete characterization of three murine models at different stages representative of Alzheimer’s disease (AD) (i.e., onset, progression and advanced). To identify genotype-to-phenotype relationships, we combined the cognitive assessment of these mice with histological analyses and a full transcriptional and protein quantification profiling from the hippocampus. As expected, we confirmed previously reported findings in these AD models, such as the age at which cognitive decline starts and when the presence of Aβ plaques becomes detectable (*27, 28*). We also confirmed that the most aggressive model (e.g., *App^NL-G-F^*) shows more changes in genes/proteins compared to healthy mice, although a significant number of dysregulated genes/proteins are shared between the three models, and that most of these changes take place at disease onset. Our comparison between AD progression and healthy aging revealed certain commonalities, such as the upregulation of microglial and inflammation markers. However, the observation that 18-mo. healthy mice do not show any cognitive decline indicates that, although accelerated aging occurs in AD models, there are other factors specifically associated with Aβ pathology. Comparison of transcriptional and quantitative protein profiles from the same mice revealed a clear correlation between mRNA and protein levels of the dysregulated genes. This observation thus supports the notion that most changes are indeed a consequence of expression changes. Interestingly, this correlation was much weaker in physiological aging, where the observed accumulation of proteins was decoupled from transcriptional changes. We also found a few proteins whose abundance increased with AD progression, while the corresponding transcript levels remained stable. These were potential cases of protein accumulation in disease conditions, and we showed that at least two of these proteins, namely lfit3 and Syt11, co-localize with Aβ plaques in the brain. Moreover, we also observed that these proteins tend to have a shorter lifespan than those found to accumulate in healthy old mice, suggesting different mechanisms of homeostasis regulation. Finally, we took advantage of the transcriptional profiles of two knock-in mice (e.g., *App^NL-F^* and *App^NL-G-F^*) to derive specific Aβ-related AD signatures, which showed a clear mobilization of microglia and astrocytes, thus reflecting the activation of the brain immune system in response to Aβ accumulation. Additionally, these effects were accompanied by the downregulation of synapse transmission processes. Interestingly, most of the functional processes associated with the characteristic AD signatures were also dysregulated in the 3xTg-AD model, thus supporting their general validity beyond AD models harboring single *App* mutations.

Despite the many efforts and a great number of clinical trials, only 5 drugs have been approved for the treatment of AD (i.e. 4 cholinesterase inhibitors and a N-methyl-D-aspartate (NMDA) antagonist with neuroprotective properties), and these are mostly symptomatic drugs that do not tackle the etiology of the disease (*17*). A lack of appropriate biomarkers, incomplete preclinical data and difficulties to start treatments at early stages of the disease may explain these failures (*19, 20*). Additionally, given the robustness of biological systems, it is also clear that the inhibition of a single target (i.e., the β-secretase) is not enough to alter the progression of the disease, and we need to look beyond the ‘one disease, one target, one drug’ paradigm. In an attempt to tackle AD from a global perspective, we profited from our recently derived compound bioactivity descriptors to find small molecules able to neutralize the changes induced by the disease (*25*). Of the ∼1M compounds, we selected six drugs, including three non-steroidal anti-inflammatory drugs (NSAIDs) and three anti-hypertensives with very different reported mechanisms of action, that showed a good potential to revert the AD signatures *in silico*. After a four-week treatment, we demonstrated that four of the six drugs, two NSAIDs (dexketoprofen and etodolac) and two anti-hypertensives (penbutolol and bendroflumethiazide) reduced the cognitive impairment in AD mice. We also demonstrated that all four compounds partially reverted the expression levels of those genes upregulated in our AD signatures, although only penbutolol was able to significantly restore the global expression levels of genes repressed in AD. Reassuringly, the hippocampi of mice treated with this antagonist of β-adrenergic receptors showed a reduction of Aβ plaques and a clear dose-dependent inhibition of Aβ_40_ and Aβ_42_ production *in vitro*.

Epidemiological studies identified NSAIDs and anti-hypertensives as protective agents against AD (*67, 68*), with the potential to target the Aβ processing pathway (*73*). Unfortunately, randomized clinical trials did not show positive effects for a number of NSAIDs (e.g., R-flurbiprofen, indomethacin, rofecoxib, naproxen or celecoxib). However, it has been reported that a 3-month treatment of a mouse model of AD with ibuprofen prevents memory impairment, although without any perceivable change in Aβ accumulation or inflammation (*66*). Recent studies also showed that fenamate NSAID has the potential to block AD pathology in animal models through COX-2-independent inhibition of the NLRP3 inflammasome (*65*). On the other hand, as hypertension in midlife is a risk factor for dementia (*74*), anti-hypertensive drugs have been proposed as potential preventive treatments for AD (*75*). Indeed, compounds targeting noradrenergic signaling, such as prazosin (*76*) and carvedilol (*77*), have shown beneficial effects on AD mouse models by alternative mechanisms, including the blockade of Aβ production and neuronal protection. Overall, despite the inconclusive and contradictory results reported, in the light of our findings we believe that NSAID and anti-hypertensive drugs might still have an opportunity as anti-AD agents.

## Methods

### Cells

SH-SY5Y cells (obtained from Jens Lüders’ lab, IRB Barcelona) were cultured in DMEM/F12 (1:1) medium supplemented with 10% FBS, glutamine and antibiotics (Thermo Fisher Scientific). 7PA2 cells overexpressing mutated APP were obtained from Dennis Selkoe’s lab (Harvard Medical School, Boston). 7PA2 cells were maintained in DMEM medium supplemented with 10% FBS, glutamine and antibiotics. CRISPR/Cas9 edited *PSEN1^M146V/M146V^* cells have been described before (*25*). For RNA interference, we used pLKO.1-puro Mission shRNA vectors (Sigma-Aldrich). Target sequences are provided in Table S7. Viral vectors were generated in 293T cells to infect SH-SY5Y cells that were selected with 2 µg/ml puromycin (Sigma-Aldrich), following the manufacturer’s recommendations. For the assays on SH-SY5Y cells, 5×10^4^ cells were initially differentiated for 3 days in neurobasal medium supplemented with B27, glutamax (Thermo Fisher Scientific), 10 µM retinoic acid (Sigma-Aldrich) and 50 ng/mL Brain-Derived Neurotrophic Factor (BDNF; Peprotech). The medium was then renewed in the presence of the indicated concentration of drugs. All drugs were dissolved in DMSO, and controls were treated with DMSO in parallel. After 3 days, supernatants were stored at −80°C for Aβ measurement, and cells were incubated for 1 h in the presence of 3-(4,5-dimethylthiazol-2-yl)-2,5-diphenyltetrazolium bromide (MTT) and then lysed in DMSO to check viability. For the assays on 7PA2 cells, 1.4×10^5^ cells were seeded in 24-well plates. The day after, medium was replaced with the indicated concentration of compounds in DMEM without FBS supplement. Cells were incubated for further 24 h before medium was collected and stored at −80°C until Aβ quantification.

### Aβ quantification

Aβ peptides were quantified with the Human β Amyloid(1–40) ELISA Kit Wako, Human β Amyloid(1–42) ELISA Kit Wako and the Human β Amyloid(1–42) ELISA Kit, High-Sensitive (FUJIFILM Wako Pure Chemical Corporation), following the manufacturer’s instructions.

### Proximity Ligation Assay

Duolink Proximity Ligation Assay (PLA) reagents were purchased from Sigma-Aldrich. Differentiated SH-SY5Y cells grown in slides were fixed and permeabilized with 0.3% Triton, 2% BSA in TBS-T (all from Sigma-Aldrich). The protocol recommended by the manufacturer was followed. Primary antibodies anti-Aβ(1–16) (Clone 6E10, 1:100, #803001, Biolengend) and anti-(human)SYT11 (1:100, ab204589, Abcam) were used. Specificity of the anti-SYT11 antibody was assessed by western blot (Fig. S6c). Images were acquired using a Zeiss LSM 780 Upright confocal, multiphoton FLIM system. For quantification, the positive dots were measured using Fiji ImageJ software. The number of cells was manually evaluated and results were expressed as the number of dots per cell.

### Drugs/compounds

BACE inhibitor AZD3839, etodolac, fenoprofen and pargyline were supplied by Selleckchem. Dexketoprofen and penbutolol were from Medchemexpress. Bendroflumethiazide was obtained from Sigma-Aldrich.

### Animals and drug treatments

The homozygous 3xTg-AD mice and the corresponding wild-type mice (C57BL/6×129/Sv mixed background) were obtained from The Jackson Laboratory. Heterozygous *App^wt/NL-F^* and *App^wt/NL-G-F^* mice were obtained from the RIKEN Brain Science Institute and used to start colonies of homozygous *App^NL-F^* and *App^NL-G-F^* mice and their respective wild-type counterparts. Mice were maintained at the animal facility of the Barcelona Institute of Science and Technology, Barcelona, Spain. Only female animals were used in the study, as several lines of evidence suggest that pathology is stronger in 3xTg-AD female mice (https://www.jax.org/strain/004807). To allow comparison between models, only female mice were used in all the models.

Mice were maintained under standard housing conditions on a 12 h light/dark cycle, with water and food ad libitum. Genotyping was done on genomic DNA from the tail or ear, using standard PCR amplification and gel electrophoresis. The DNA primers (Sigma-Aldrich) used for genotyping are listed in Table S7.

During the treatment period, *App^NL-G-F^* mice received a daily intraperitoneal injection of the indicated drugs for 4 weeks. Etodolac (50 mg/kg), dexketoprofen (15 mg/kg), fenoprofen (50 mg/kg) and bendroflumethiazide (20 mg/kg) were dissolved in a mixture of 5%Tween-80 (Sigma-Aldrich), 25% propylene glycol (Sigma-Aldrich), 25% Poly(ethylene glycol) (Sigma-Aldrich) and 45% phosphate-buffered saline (PBS). Penbutolol (20 mg/kg) and pargyline (20 mg/kg) were dissolved in PBS (veh.2). The drug doses were chosen according to previous publications on these compounds or similar compounds (*78–83*). Groups of *App^wt^* and *App^NL-G-F^* mice were also treated with the vehicles. The Ministry of Agriculture, Livestock, Fisheries and Food, Government of Catalonia, Spain, authorized all the mouse experiments (authorization numbers 8478, 8527 and 10382).

### Cognitive tests

The Novel Object Recognition (NOR) test was performed following the protocol described in (*29*). On the first day, the habituation session, mice were placed in the empty box and allowed to explore for 5 min. Twenty-four hours after habituation, mice were placed in the same box in the presence of two identical objects (familiarization session) and allowed to explore for 10 min. The pair of objects was randomized and counterbalanced between mice. Twenty-four hours later (test session), mice were placed in the same box, but a novel object substituted one of the known objects, and the mice were allowed to explore for 5 or 10 min. The position of the novel object (left or right) was randomized and counterbalanced between mice. The objects and the box were cleaned with odorless soap between trials. All sessions were recorded. Object exploration was defined as the mouse sniffing or touching the object with the nose while looking at it. Climbing onto the object or chewing was not considered exploration. An experimenter blinded to the mouse condition tested assessed the exploratory activity manually. The amount of time exploring the novel object was expressed as a percentage of the total exploration time. Ability to discriminate the novel object was evaluated by performing a one-sample t-test against a hypothetical value of 50%.

The Y-maze test was conducted as described in (*84*). The Y-maze apparatus consisted of three identical arms set at angles of 120°. Each animal was placed at the center of the maze and was allowed to explore it freely for 5 min. The maze was cleaned with odorless soap between trials. All sessions were recorded. The four paws of the mouse had to enter an arm to count as an arm entry. Alternation behavior was defined as consecutive entries into each of the three arms without repetition (ABC, CBA, etc.). We expressed the results as the percentage of spontaneous alternations, which was calculated by dividing the number of alternations by the total number of possible alternations (total arm entries-2) X 100.

Additional analyses of the videos were performed with dedicated tracking software (SMART video tracking, Panlab). The results obtained were in concordance with our observations. However, the inability of the software to discriminate between mouse head and tail in all the situations led us to rely on manual analysis for a more accurate result.

### Mouse brain tissue preparation

Mice were anesthetized with an intraperitoneal (i.p.) mixture of ketamine/xylazine (100/10 mg/kg) and perfused intracardially with 0.9% saline solution. Brains were removed and cut in two hemispheres. The left hemisphere was snap-frozen in cold isopentane for 1 min, saved in dry ice and stored at −80°C for immunofluorescence assays. The right hemisphere was dissected into the hippocampus and cortex, snap-frozen in dry ice and stored at −80°C. Only hippocampal tissue was used for RNA and protein extraction.

### Immunofluorescence staining

20-µm coronal sections of the entire brain were cut using a Leica CM1900 cryostat. Sections were mounted on SuperFrost Ultra Plus slides (TermoFisher) and fixed in cold acetone for 10 min at room temperature (RT). After drying at RT, the mounted slides were saved at −20°C until further processing. Brain sections were washed in PBS for 5 min and incubated with blocking buffer (1% BSA, 5% goat serum and 0.2% Triton in PBS) for 20 min at RT in a humidifier chamber. After the slides had been washed in PBS, the sections were incubated overnight at 4°C with the primary antibodies diluted in PBS with 1% BSA and 0.2% Triton. The following primary antibodies were used: anti-Aβ (clone D54D2, 1:100, #8243, Cell Signaling), anti-human PHF-tau (clone AT-8, 1:100, #90206, Innogenetics), anti-Aβ(1–16) (Clone 6E10, 1:100, #803001, Biolengend), anti-Syt11 (1:100, #270003, Synaptic Systems) or anti-Ifit3/P60 (1:50, ab76818, Abcam). After PBS washes, sections were incubated for 1 h at RT with the corresponding goat anti-rabbit Alexa Fluor 488 (1:250, ThermoFisher), donkey anti-rabbit Alexa Fluor 647 (1:250, ThermoFisher) or goat anti-mouse Alexa Fluor 568 (1:250, ThermoFisher) diluted in PBS with 1% BSA and 0.2% Triton. Sections were counterstained with 2 µg/ml Hoechst for 5 min at RT, washed in PBS and finally coverslipped using Fluoromount-G (EMS). Control sections (without primary antibody) were used to differentiate specific from non-specific staining. Images were acquired using a Nikon Eclipse E800M microscope equipped with an Olympus DP72 camera or a Zeiss LSM 780 Upright confocal, multiphoton FLIM system. For quantification, two coronal sections per mouse were analyzed, and the immunoreactive areas were measured using Fiji ImageJ software.

### RNA extraction, mRNA library preparation and sequencing

Total RNA was extracted from mouse hippocampal samples using a RNeasy Mini kit (Qiagen), following the manufacturer’s protocol, and sent for whole transcriptome sequencing at the Centro Nacional de Análisis Genómico (CNAG-CRG). Total RNA was assayed for quantity and quality using the Qubit RNA BR Assay kit (Thermo Fisher Scientific) and RNA 6000 Nano Assay on a Bioanalyzer 2100 (Agilent).

The RNA-Seq libraries were prepared from total RNA using KAPA Stranded mRNA-Seq Kit Illumina Platforms (Roche-Kapa Biosystems) with minor modifications. Briefly, after poly-A based mRNA enrichment with oligo-dT magnetic beads and 500 ng of total RNA as the input material, the mRNA was fragmented (resulting RNA fragment size was 80-250 nt, with the major peak at 130 nt). First-strand cDNA was synthesized using random priming. Second strand cDNA was synthesized in the presence of dUTP instead of dTTP, to achieve strand specificity. The blunt-ended double-stranded cDNA was 3’adenylated and Illumina indexed adapters (Illumina) were ligated. The ligation product was enriched with 15 PCR cycles and the final library was validated on an Agilent 2100 Bioanalyzer with the DNA 7500 assay.

Each library was paired-end sequenced using TruSeq SBS Kit v4-HS, with a read length of 2×76bp. On average, we generated 53 million paired-end reads for each of the 48 samples of *App^NL-G-F^*, *App^NL-G-F^*, *App^wt^* mice, and 80 million reads for each of the 24 samples of 3xTg-AD and C57BL/6×129/Sv mice in a fraction of a sequencing lane on HiSeq2000 (Illumina, Inc) following the manufacturer’s protocol. Image analysis, base calling and quality scoring of the run were processed using the manufacturer’s software Real-Time Analysis (RTA 1.18.64) and followed by generation of FASTQ sequence files by CASAVA.

### RNA-Seq data analysis

STAR software (Dobin et al., 2013) was used to align the raw RNAseq reads to the mouse reference genome (GRCm38/mm10 and GENCODE vM15 genome annotations (*85*)). MAPT, PSEN1, and humanized APP and/or mutated sequences were included according to the genotype of the mouse model analyzed. We used casper (*86*) to quantify the expression of all transcript isoforms, which were aggregated at gene level and quantile-normalized. To reduce biases caused by low expression, we considered only genes with at least 0.2 RPKMs in 90% of the samples. This filter was applied separately to the 48 samples of APP knock-in mice and their controls (GSE168430), to the 24 samples of 3xTg mice and their controls (GSE168428), and to the 32 samples of drug-or vehicle-treated mice (GSE168429). We used the Limma package (*87*) to perform differential expression analysis based on empirical Bayes moderated t-statistics, including experimental batch as covariate. We performed all pairwise comparisons along the genotype- and time-axes to describe AD progression and physiological aging. To derive the *AD signatures*, we specified an additional model to analyze the APP knock-in dataset, in which we considered age as a continuous variable together with its interaction with genotype. These models were used to estimate the linear association of gene expression with age for each genotype. Multiple comparisons correction was performed using the Benjamini-Hochberg algorithm. For visualization purposes, quantile-normalized expression values were corrected by batch using the removeBatchEffect function from Limma. Genes were ranked by multiplying their fold-change sign by the -log10 (p-value) for pre-ranked GSEA (*88*). We used the prcomp function in R to perform a principal components analysis (PCA) of the gene expression matrices before and after batch adjustment. We used the pvca Bioconductor package to perform a principal variance component analysis (PVCA) aimed at identifying the most prominent sources of variability n each dataset.

### Mass spectrometry sample preparation

Frozen mouse hippocampi were homogenized in 0.2 ml of pre-cooled homogenization buffer (0.3 M sucrose, 10 mM MOPSNaOH and 1mM EDTA) supplemented with complete protease inhibitor cocktail (Roche). 50 µl of lysis buffer (8% SDS and 0.2 M DTT in 0.2 M Tris-HCl pH 7.6) was added to 50 µl homogenate and the mixture was incubated for 3 min at 95°C. Once cooled, the total protein extracts were stored at −20°C until quantification. Protein samples were quantified using the Pierce 660 Protein Assay Kit, reduced with tris (2-carboxyethyl) phosphine (TCEP), alkylated, and digested with trypsin. After digestion, all samples were isotopically labeled with the corresponding iTRAQ-8plex reagent according to the experimental design and following the manufacturer’s instructions (Thermo Fisher Scientific). iTRAQ labels were randomized, minimizing the co-occurrence of labels within a biological condition. Samples were desalted using C18 and strong cationic exchange tips. Each batch was fractionated off-line by basic reversed-phase chromatography. Sample fractionation was performed with a Zorbax 300 Extend-C18 column (2.1 x 150 mm, 3.5 mm) in an AKTA micro ETTAN gradient LC system (Amersham Biosciences).

Peptides were separated in a total of 84 collected fractions and grouped into 24 fraction groups per batch and dried via vacuum centrifugation. Fraction groups (∼5µg) were reconstituted in 48µl of 3% acetonitrile (ACN), 1% formic acid (FA) aqueous solution for nano LC-MS/MS analysis.

### Mass spectrometry analysis

Mass spectrometry data were collected on an Orbitrap Fusion Lumos™ Tribrid mass spectrometer (Thermo Scientific) equipped with a Thermo Scientific Dionex Ultimate 3000 ultrahigh pressure chromatographic system (Thermo Fisher Scientific) and an Advion TriVersa NanoMate (Advion Inc. Biosciences) as the nanospray interface. Peptide mixtures (6µl) were loaded into a µ-Precolumn (300µm i.d x 5 mm, C18 PepMap100, 5 µm, 100 Å, C18 Trap column; Thermo Fisher Scientific) at a flow rate of 15 µL/min and separated using a C18 analytical column (Acclaim PepMap TM RSLC: 75 µm x 75 cm, C18 2 m, nanoViper) with a flow rate of 200 nl/min and a 210 min run, comprising three consecutive steps with linear gradients from 1 to 35% B in 180 min, from 35 to 50% B in 5 min, and from 50 % to 85 % B in 2 min, followed by isocratic elution at 85 % B in 5 min and stabilization to initial conditions (A= 0.1% FA in water, B= 0.1% FA in ACN) The mass spectrometer was operated in a data-dependent acquisition (DDA) mode. In each data collection cycle, one full MS scan (400-1600 m/z) was acquired in the Orbitrap (120k resolution setting and automatic gain control (AGC) of 2 x 10^5^). The following MS2-MS3 analysis was conducted with a top-speed approach. The most abundant ions were selected for fragmentation by collision-induced dissociation (CID). CID was performed with collision energy of 35%, 0.25 activation Q, an AGC target of 1 x 10^4^, an isolation window of 0.7 Da, a maximum ion accumulation time of 50 ms and turbo ion scan rate. Previously analyzed precursor ions were dynamically excluded for 30 s. For the MS3 analyses for iTRAQ quantification, multiple fragment ions from the previous MS2 scan (SPS ions) were co-selected and fragmented by HCD using a 65% collision energy and a precursor isolation window of 2 Da. Reporter ions were detected using the Orbitrap with a resolution of 30k, an AGC of 1 x 10^5^ and a maximum ion accumulation time of 120 ms. Spray voltage in the NanoMate source was set to 1.60 kV. RF Lens were tuned to 30%. The minimal signal required to trigger MS to MS/MS switch was set to 5,000. The mass spectrometer was working in positive polarity mode and single charge state precursors were rejected for fragmentation.

Database searches were performed with Proteome Discoverer v2.1.0.81 software (Thermo Fisher Scientific) using Sequest HT search engine and SwissProt Mouse_canonical_2016_11 including contaminants and MAPT, PSEN1 and APP humanized and/or mutated sequences according to each of the mouse models analyzed. Search was run against targeted and decoy database to determine the false discovery rate (FDR). Search parameters included trypsin, allowing for two missed cleavage sites, carbamidomethyl in cysteine and iTRAQ 8-plex peptide N-terminus as static modification and iTRAQ 8plex in K/Y, methionine oxidation and acetylation in protein N-terminus as dynamic modifications. Peptide mass tolerance was set to 10 ppm and the MS/MS tolerance to 0.6 Da. Peptides with an FDR < 1% were considered as positive identifications with a high confidence level. The mass spectrometry proteomics data have been deposited to the ProteomeXchange Consortium via the PRIDE (*32*) partner repository with the dataset identifier PXD024538.

All computations were performed in the R statistics framework. The iTRAQ reporter intensities were filtered by contamination flag set to “False”, average reporter signal to noise ratio larger than 3, intensity value larger than 1000, and the Peptide Quan Usage flag set to “Use”. For peptides without a unique assigned protein, the protein with maximum total intensity was defined as master protein. Filtered intensities were normalized within each processing batch by a size factor computed as in (*89*). To do this, intensities were summarized via the median for each protein and sample. A reference sample was computed as the mean value of all samples for each protein. Size factors were computed as the median ratio of each protein against the reference sample. Once the size factors were obtained, all PMSs were divided by the corresponding factor and the log2 value (after adding 1) was computed.

For each protein, a linear model was fitted with or without random effects depending on the number and combination of peptides measured in each age*genotype group. When more than one peptide was measured in more than one group and throughout several fractions, peptide, fraction (within peptide), and sample were included in the model as random effects, with age*genotype and batch as fixed effects. When only one peptide or one fraction was found, random effects were dropped accordingly. When no repeated measures existed for a given protein, a linear model was fitted with the same fixed effects. Mixed effects models were fitted using the lmer function from the lme4 package (*90*), while linear models were fitted with the lm function of the stats package. To perform the desired contrasts, the glht function from the multcomp package was used without p-value adjustment (*91*). Multiple comparisons correction was performed a posteriori for each contrast using the p.adjust function with the Benjamini-Hochberg method.

For visualization purposes, normalized intensities were corrected by peptide, sample and batch using the model coefficients.

### Statistics

Data were analyzed with the Prism statistical package. Unless otherwise indicated in the figure legend, P-values were calculated using an unpaired, one-tailed, Student t-test. To control possible batch effects in transcriptomics and proteomics, we used a blocked/randomized design. Biological replicates were confounded with the processing batches (4 batches in each mouse model). iTRAQ labels were randomized to minimize the co-occurrence of labels within a biological condition.

### Virtual signature-based screening of compounds

We devised a strategy to select the most promising small molecules to modify the biology of AD within the universe of the ∼1M bioactive compounds catalogued in the CC. The CC aggregates 25 data types for the molecules, organized in five levels (A-E) and sublevels (1–5). On the one hand, we defined three ‘pharmacological queries’ to identify compounds that were (1) chemically similar (P < 0.001 in the A1-5 bioactivity spaces of the CC) to drugs that have entered clinical trials against AD, or that shared a significant number of targets with them (B4 signatures). More specifically, we considered drugs thought to target amyloid fibrils and plaques, tau, cholesterol, inflammation, neurotransmitters and cholinergic receptors (*62*). We also (2) selected small molecules that showed similar transcriptional (D1) or cell sensitivity (D2) to these drugs. In addition, (3) we looked for small molecules known to bind putative AD targets, as defined by OpenTargets (*63*). Furthermore, we designed two ‘biological queries’ to capture connectivities between the discovered molecular changes in AD models and the bioactivity data available in the CC. In particular, we looked for (4) compounds that trigger transcriptional responses able to neutralize the changes observed in the AD mice in the CC D1 space. Finally, (5) we used the LINCS L1000 resource (*92*) to find perturbation experiments, mainly shRNA, that could also revert our AD transcriptional signatures, used them as putative AD targets and looked for compounds in the CC able to inhibit their activity (B spaces). Molecules selected in at least one of the 5 queries were kept; these candidates based on virtual screening can be found in Table S5, together with detailed scores for each of the queries. Scores are represented as empirical -log10 p-values obtained over the CC universe of >800k molecules.

Additional calculations were done to facilitate the selection of compounds for experimental screening. In particular, we trained machine-learning classifiers based on the BBBP and BACE datasets available from MoleculeNet (*93*), and Aβ40, Aβ42 and Aβ40/Aβ42 ratio from a previous compound screening performed in iPSC cells (*94*) (binarization cut-offs: Aβ 0.8, ratio 0.9). As a machine-learning method, we used an ensemble-based approach (extra-trees classifiers) and, as features, we used CC signatures. Ensemble-based methods applied to CC signatures have shown exceptional performance across a wide range of benchmarking tasks (*95*). In a stratified 5-fold cross-validation, we obtained ROC AUC > 0.918, 0.874, 0.774, 0.690 and 0.874 for BBP, BACE, Aβ40, Aβ42 and Aβ ratio models, respectively.

## Acknowledgements

We would like to thank Prof. E Wanker and his team at MDC-Berlin for helpful discussions. The authors acknowledge the support of the IRB Barcelona Proteomics and Biostatistics Units. PA acknowledges the support of the Generalitat de Catalunya (RIS3CAT Emergents CECH: 001-P-001682 and VEIS: 001-P-001647), the Spanish Ministerio de Economía y Competitividad (BIO2016-77038-R) and the European Research Council (SysPharmAD: 614944).

## Author contributions

EP, SB, LM, MD-F and PA designed the study. EP, SB and VA performed all the *in vivo* and *in vitro* studies. EP, LM and TJ-B analyzed the results, with the help of AB-Ll and CS-OA to process and analyze the transcriptomics and proteomics data. MD-F implemented the computational strategy to find potential chemical modulators of AD. MG and EdO performed quantitative proteomics experiments. TCS and TS provided the knock-in mouse models. All the authors discussed and interpreted the results. EP, LM, MD-F and PA wrote the manuscript, which has been approved by all the authors.

## Data and materials availability

All the mRNAseq data have been deposited in the Gene Expression Omnibus (GEO; (*31*) database with the identifier GSE168431. The mass spectrometry proteomics data have been deposited to the ProteomeXchange Consortium via the PRIDE (*32*) partner repository with the dataset identifier PXD024538.

## Supplementary Figures and Tables

**Figure S1.**
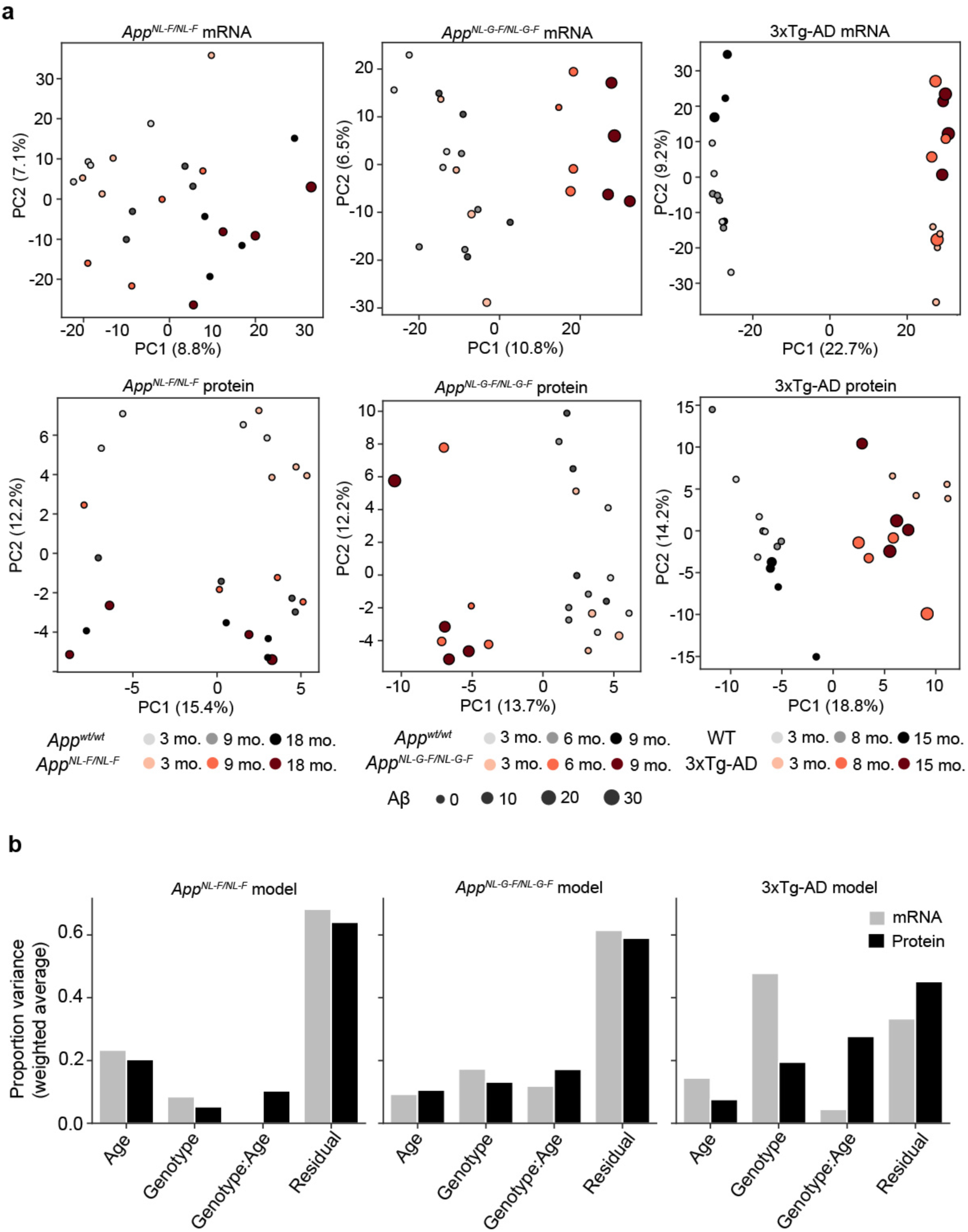
Principal Component Analysis (PCA) of hippocampus gene expression influenced by age and genotype. **(a)** PCA plot showing distribution of samples corresponding to the *App^NL-F^* and *App^NL-G-F^* models and their respective *App^wt^* controls (gray colors) at different ages (left panel). On the right panel, the PCA plot shows the separation for 3xTg-AD mice and the corresponding wild-type controls (C57BL6/129SvJ; gray colours). Size of the dot is proportional to the quantification of Aβ for each of the samples. **(b)** Principal Variance Component Analysis (PVCA) showing which sources of variability are most prominent in each dataset after adjustment for batch (mRNA) or random effects (protein).

**Figure S2.**
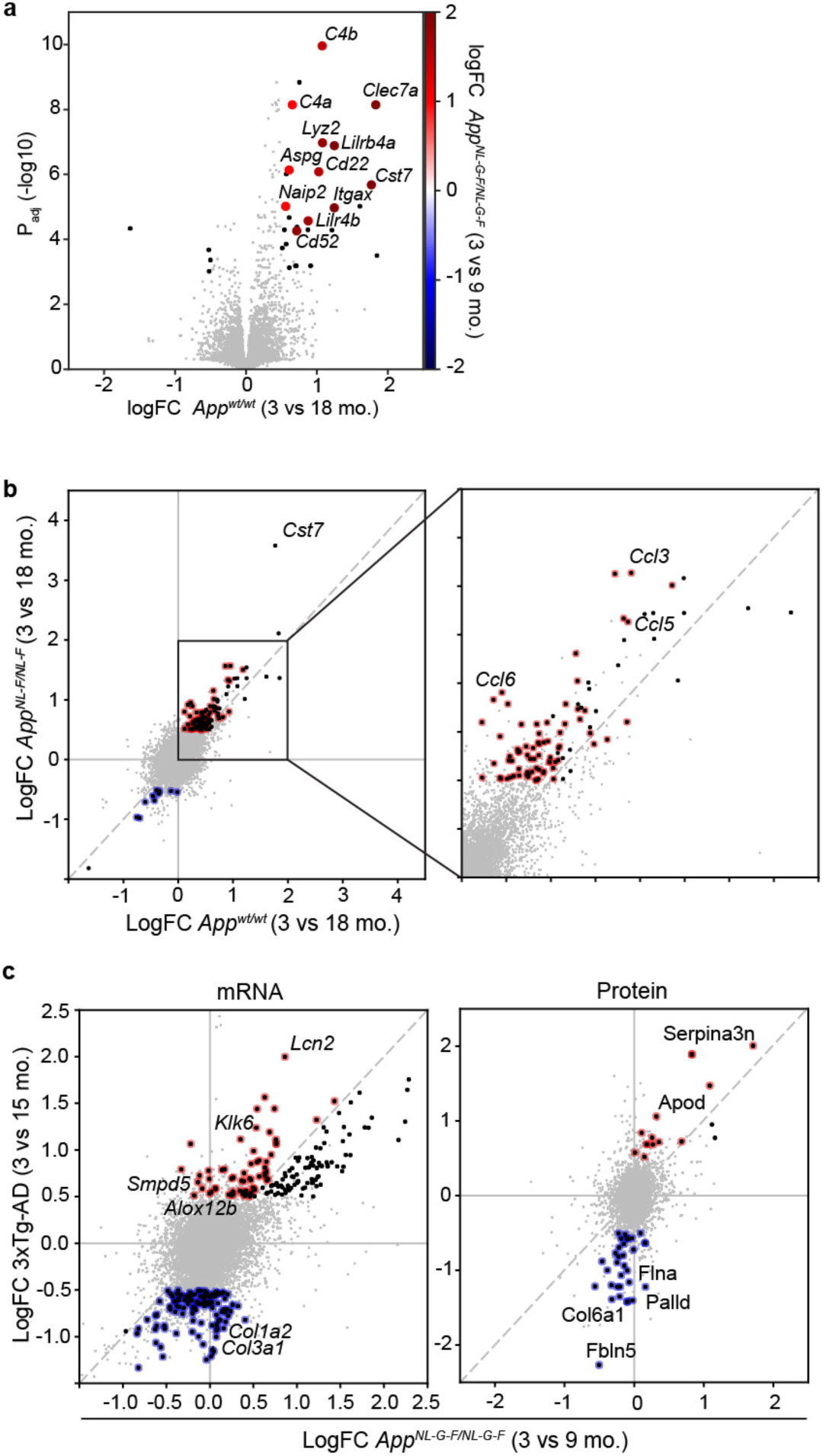
**(a)** A Volcano plot reporting the gene expression logarithmic fold-change (logFC; X-axis) and the adjusted p-value (*P*adj; Y-axis) when analyzing the 3- vs. 18-month-old (mo.) *App^wt^* mice comparison (n=4). Genes that show a higher fold-change in the 3- vs. 9-mo. *App^NL-G- F^* mice comparison are highlighted using a color gradient. **(b)** Scatter plot representing fold-changes of the 3- vs. 18-mo. *App^wt^* mice (X-axis) compared with the fold-changes in the 3- vs. 18-mo. *App^NL-F^* mice (Y-axis). Black dots are used to represent significantly up-/down-regulated genes in the *App^NL-F^* model (Abs. logFC>0.5, FDR < 5%), while colored dots identify genes significantly up- (red) or down-regulated (blue) only in the *App^NL-F^* comparison but not in the *App^wt^* comparison (n=4). **(c)** Scatter plot comparing Log fold-changes of genes (left panel) and proteins (right panel) of *App^NL-G-F^* mice comparing 3- vs. 9-mo. ages (X-axis) and 3xTg-AD mice comparing 3- vs. 15-mo. ages (Y-axis). Black dots are used to represent significant changes in the 3xTg-AD model (Abs. logFC>0.5, FDR<5%). Red (upregulated) and blue (downregulated) dots represent genes/proteins with an absolute logFC higher for the 3- vs. 15-mo 3xTg-AD. comparison than in the comparison of 3- vs. 9-mo. *App^NL-G-F^* mice (n=4).

**Figure S3.**
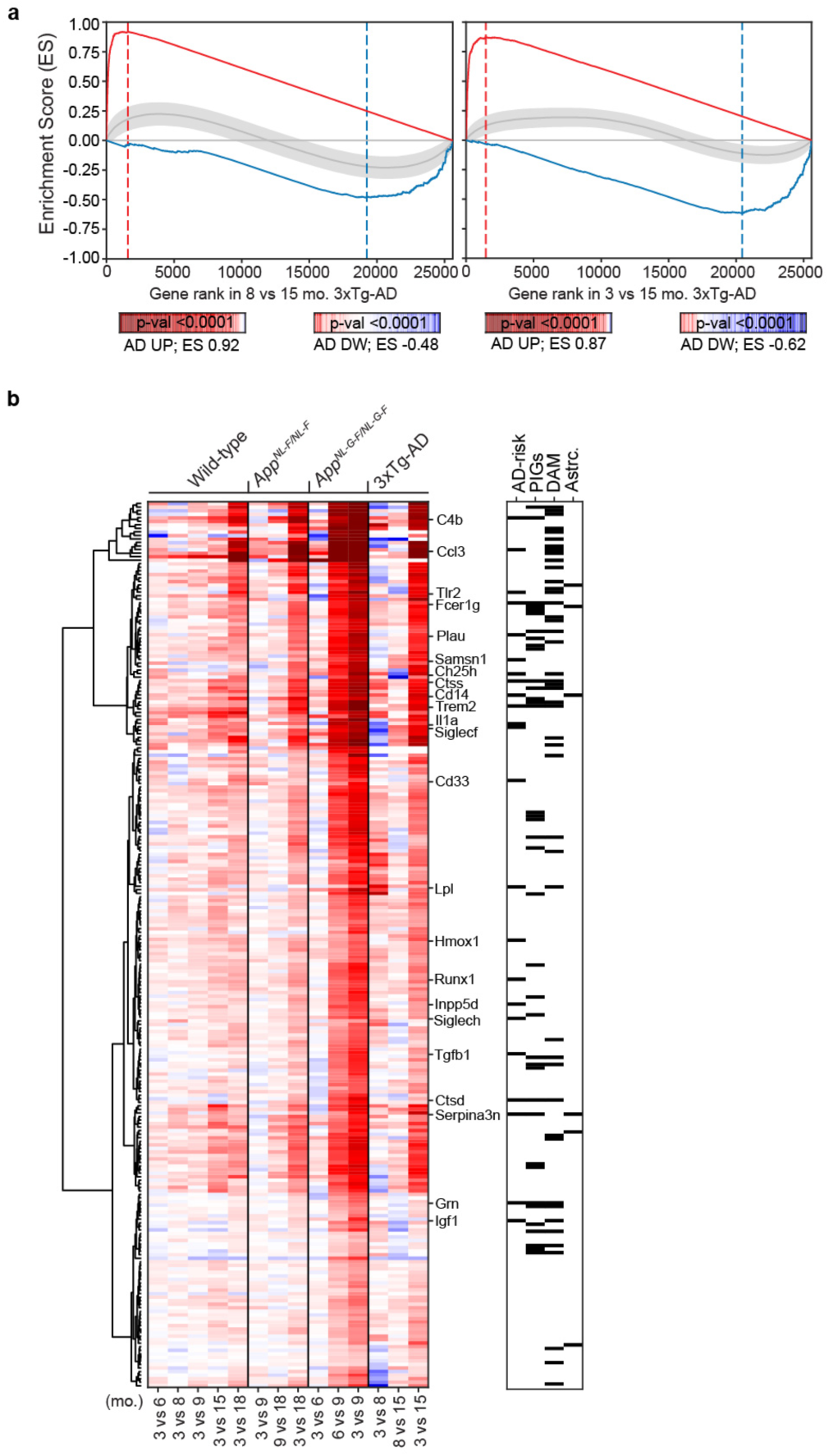
Analysis of the *AD signatures*. **(a)** Validation of the *AD signatures* in 3xTg-AD mice. In the X-axis, genes are ranked by their differential expression in 15-mo. 3xTg-AD mice compared to earlier time points (8 and 3 mo.). The Y-axis represents the running Enrichment Score (ES) computed for the *AD-UP signature* (red) and *AD-DW signature* (blue), which were derived from *App^NL-G-F^* mice and are recapitulated in the 3xTg-AD mouse model. RNAseq data were obtained from n=4 mice per condition. **(b)** Heatmap showing the progression of transcriptional changes captured by the *AD-UP signature* across models. Each gene is represented by a row of colored tiles, the color representing the expression level for the indicated condition (red, upregulated; blue, downregulated). On the right, black squares indicate whether the gene is present in a list of genes linked to AD risk (AD-risk), genes detected as proximal to amyloid plaques (Plaque-induced Genes, PIG), or in a list of genes identifying Disease-associated microglia (DAM). For the AD-risk genes, name is indicated next to the heat map.

**Figure S4.**
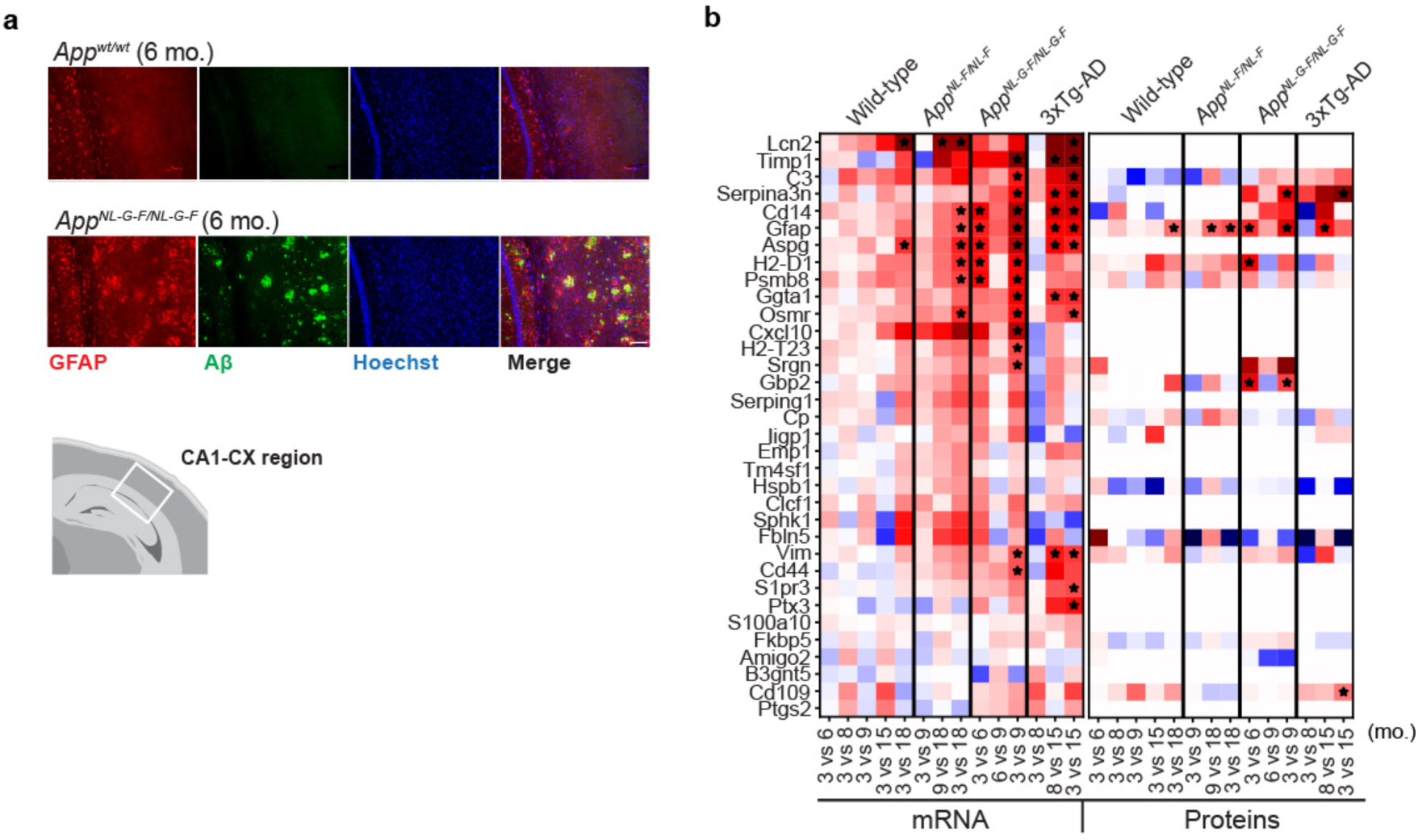
Analysis of the astrocytic signature. **(a)** Representative microphotographs of the CA1-CX region of slices of the brain of 6-mo. *App^wt^* and *App^NL-G-F^* mice stained with an anti-GFAP antibody (red), an anti-Aβ antibody (green) and Hoechst dye (blue). Scale bar represents 100 µm. N=3. **(b)** A heatmap showing the LogFC values corresponding to the gene expression and protein analysis of the 34 genes that constitute the *AD-astrocytosis* signature as defined in the main text. Each gene is represented by a row of colored tiles, the color representing the fold change in expression for the indicated condition (red, upregulated; blue, downregulated). Stars indicate statistically significant changes (|logFC|>0.5 for mRNA, |logFC|>0.25 for protein; FDR<5%).

**Figure S5.**
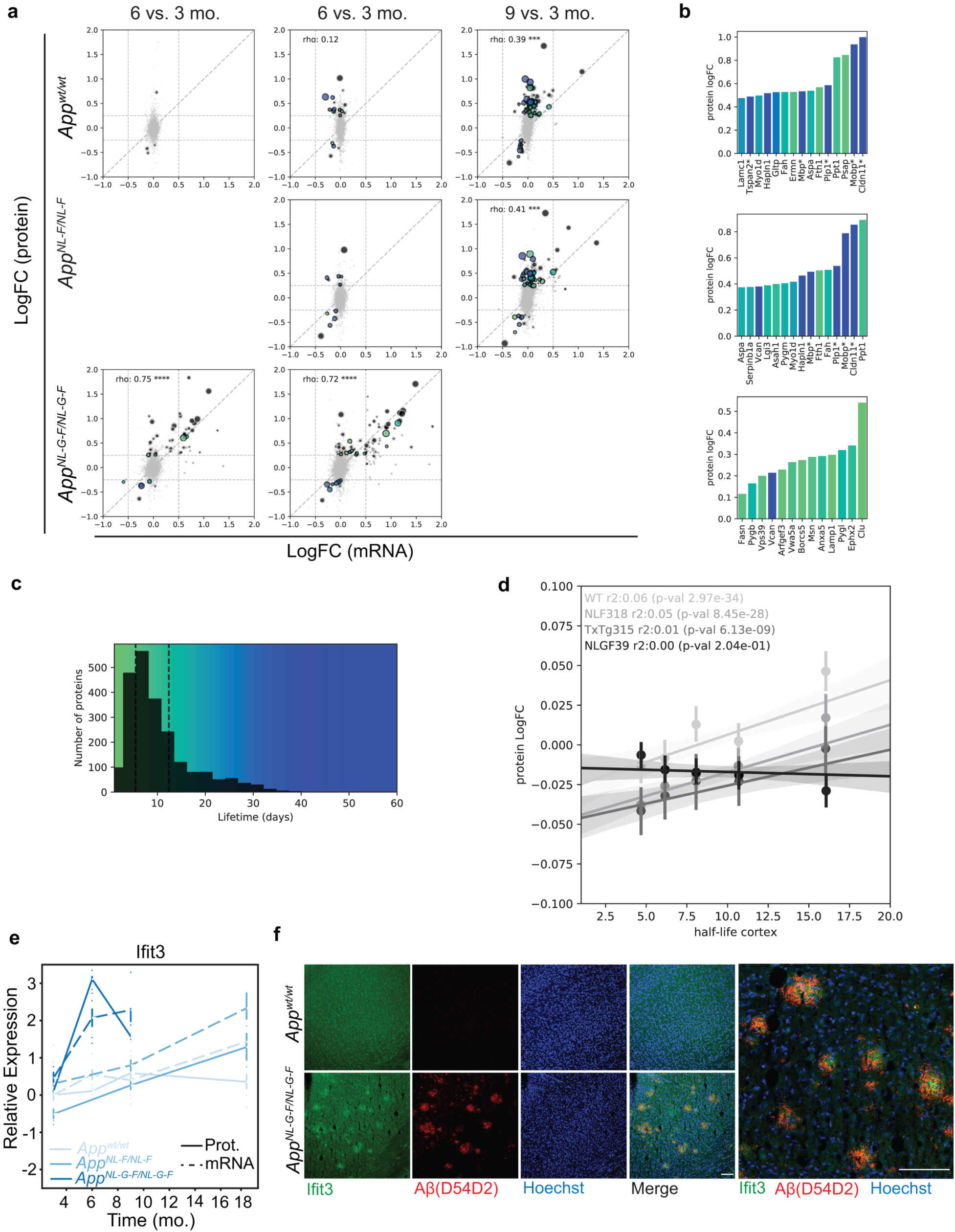
Analysis of mRNA and protein correlation in the different comparisons. **(a)** Scatter plot depicting the Logarithm of the Fold-Change (LogFC) of mRNA (X-axis) and protein (Y-axis) for the comparison of 6-, 9-, and 18-mo. with respect to 3-mo. *App^wt^*, *App^NL-F^*, and *App^NL-G-F^* mouse models. Proteins whose LogFC(protein)>0.25 and their corresponding LogFC(mRNA)>0.5 at a 5% FDR threshold are highlighted. Dot color indicates the half-life of the proteins in cortex homogenates, as defined by (doi: 10.1038/s41467-018-06519-0). Dot size is proportional to the negative logarithm of the adjusted p-value of protein changes. **(b)** Bar plot representation of the strongest discordant changes in protein abundance with estimated protein lifetimes represented as a color scale. Myelin proteins (highlighted with an asterisk) tend to accumulate in aged mice, together with other long-lived proteins. **(c)** Distribution of the estimated protein lifetime in our dataset and color scale used in **a** and **b**. Dashed lines indicate the upper and lower quartiles of the distribution (12.5 and 5.5 days, respectively). **(d)** Association between discordant changes in protein abundance with respect to protein lifetimes. The lines show the fit of a linear regression model with points indicating the average protein LogFC of 5 equally sized bins, with error bars indicating the 95% confidence interval. Proteins showing concordant changes with mRNA were excluded from this analysis. The coefficients of determination (r2) are very modest but significant in aged mice (*App^wt^*, *App^NL-F^* and 3xTg-AD) but not in younger and more aggressive AD mouse models (*App^NL-G-F^*). **(e)** Ifit3 protein (continuous line) and mRNA (dashed line) levels at different time points relative to the 3 mo. *App^wt^* model are shown for the *App^NL-G-F^* (strong blue), *App^NL-F^* (medium blue) and *App^wt^* (corresponding to the *App^NL-F^* model; light blue) mice. N=4. **(f)** Representative micrographs of brain slices of 6-mo. *App^NL-G-F^* mice stained with an anti-Ifit3 antibody (green), the anti-Aβ antibody D54D2 (red) and Hoechst dye (blue). Scale bars represent 100 µm (n=3).

**Figure S6.**
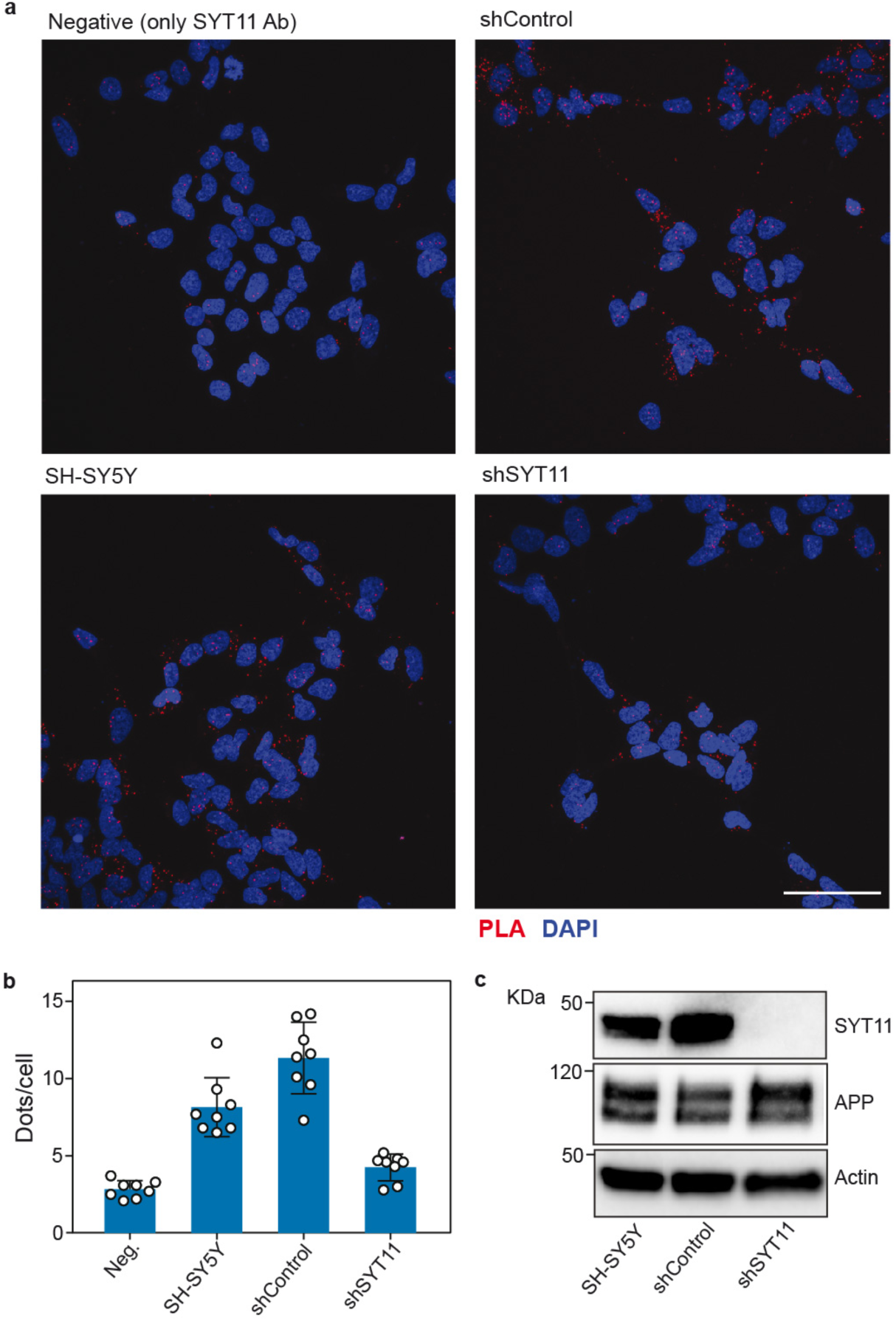
PLA analysis of SYT11 and App interaction in SH-SY5Y cells. **(a)** Representative micrographs of differentiated SH-SY5Y cells that were treated only with anti-SYT11 antibody (negative control) or both anti-SYT11 and anti-App antibody (clone 6E10; SH- SY5Y). SH-SY5Y stably expressing a control shRNA (shControl) and an shRNA targeting SYT11 (shSYT11) were also used. Bar indicates 50 µm. **(b)** Quantification of 8 different fields corresponding to the experiment shown in (a). PLA positive dots (red) were quantified and expressed as dots per number of cells. Cell nuclei are stained with DAPI (blue) to identify individual cells. Mean±SD are shown, n=8. **(c)** Western blot of differentiated, non-transduced, SH-SY5Y cells (SH-SY5Y) or cells expressing control shRNA (shControl) or a shRNA targeting SYT11 (shSYT11). In all panels, a representative experiment out of three independent experiments is shown.

**Figure S7.**
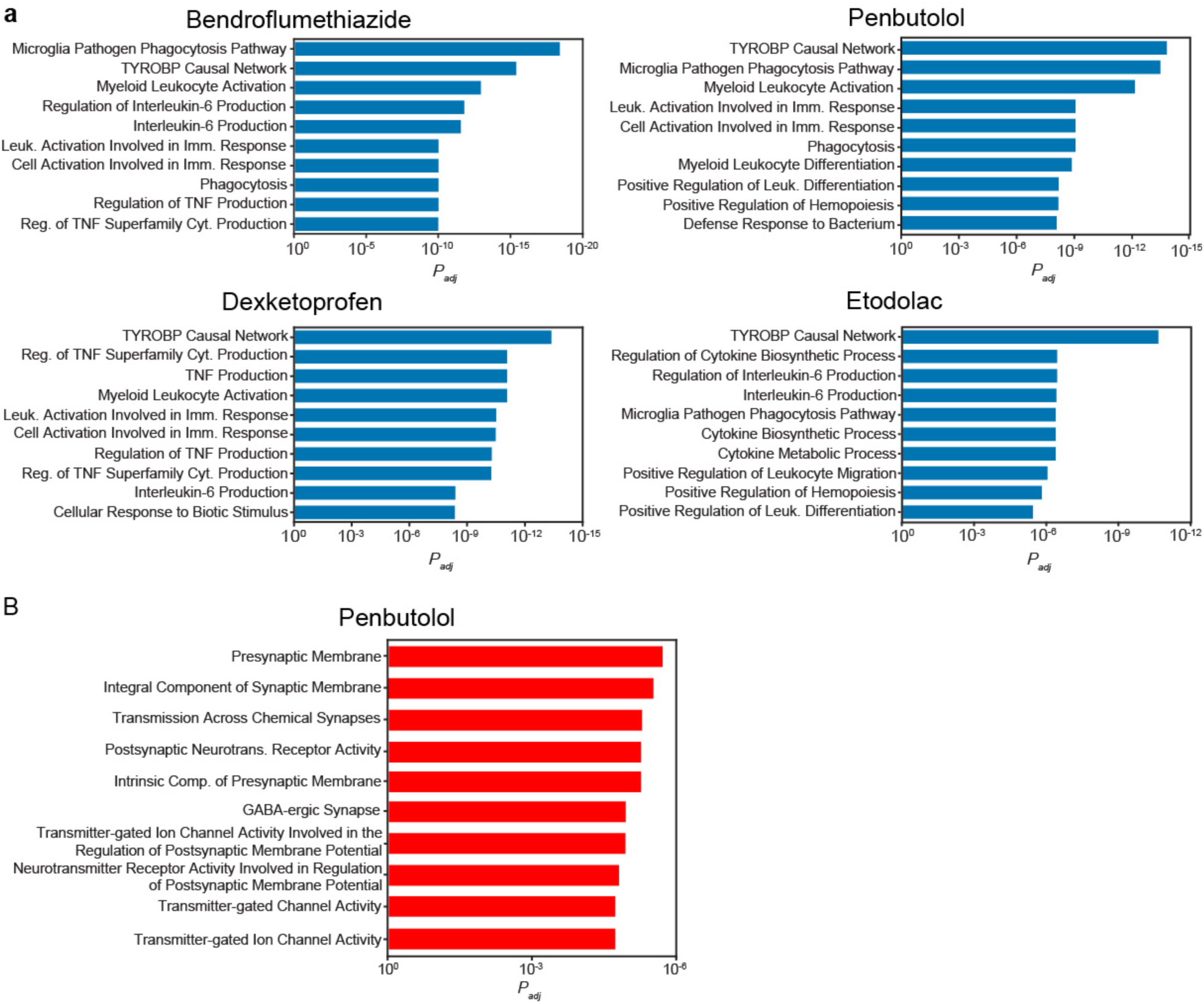
Analysis of the signature reversion in treated mice. Functional enrichment analysis of the leading-edge of the **(a)** *AD-UP signature* reversion (blue) or **(b)** *AD-DW signature* reversion (red) induced by the different treatments. The bars show the adjusted p-value of the 10 most overrepresented biological pathways. RNAseq data were obtained from n=4 mice per condition.

**Figure S8.**
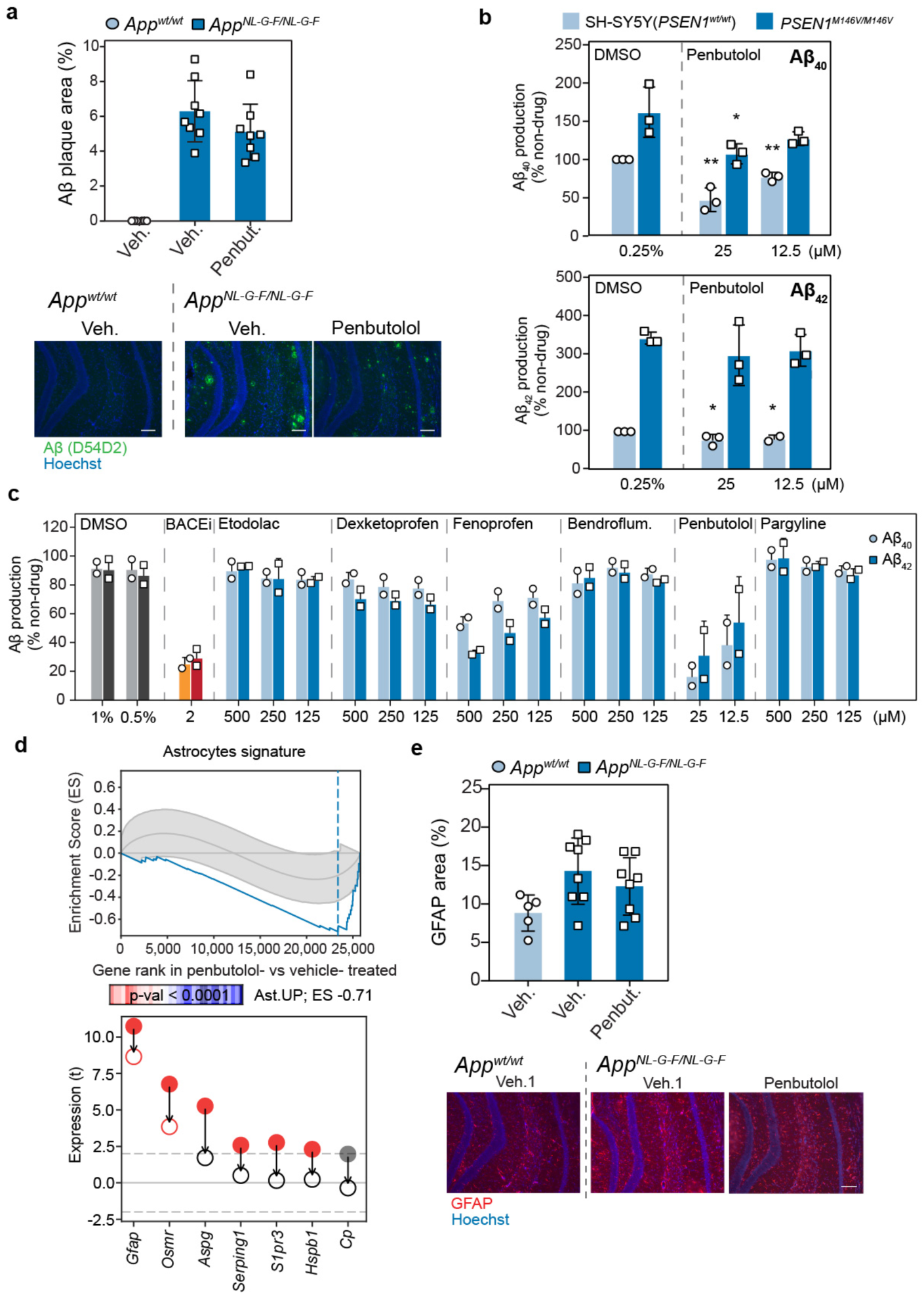
Penbutolol prevents Aβ accumulation and astrocytosis. **(a)** Percentage of Aβ- positive area measured in brain sections of 6-mo penbutolol- and vehicle-treated *App^wt^* (circles) and *App^NL-G-F^* (squares) mice. Mean±SD are shown (n= 3-4 for each condition). Below, representative microphotographs of the DG-CA1 region of the hippocampus stained with an anti-Aβ antibody (green) and Hoechst dye (blue) are shown. Scale bars represent 100 µm. **(b)** Effect of penbutolol in a cellular AD model. Normalized Aβ_40_ and Aβ_42_ secretion in differentiated wild-type (*PSEN1^wt^*) or mutated (*PSEN1^M146V/M146V^*) SH-SY5Y cells treated with the indicated concentration of penbutolol. Mean ± SD of 3 independent experiments are shown. Unpaired Student’s t-test (** P < 0.01, * P < 0.05). **(c)** Aβ production in 7PA2 cells. Normalized Aβ_40_ and Aβ_42_ secretion in 7PA2 cells treated with the indicated compounds. Percentage of production of Aβ_40_ (circle, light colors) and Aβ_42_ (square, dark colors) compared with cells treated in the absence of drug. DMSO (gray) and a BACE1 inhibitor (BACEi; AZD3839) are used as controls. Mean ± SD of two independent experiments are shown. **(d)** Reversion of genes belonging to the previously described *astrocytosis signature*. As in Figure 5, the top panel shows a graphic representation of the Enrichment Score (ES). In the X-axis, genes are ranked by their differential expression in the comparison of penbutolol- vs. vehicle-treated *App^NL-G-F^* mice. The Y-axis represents the running ES computed for the *astrocytosis signature*, which tends to be downregulated by penbutolol treatment in *App^NL-G-F^* mice. On the bottom panel, example genes that are up- (red) or down-regulated (blue) (t-score) in the vehicle-treated *App^NL-G-F^* vs. vehicle- treated *App^wt^* comparison (bold dots) or in the penbutolol-treated *App^NL-G-F^* vs. vehicle-treated *App^wt^* comparison (empty dots). RNAseq data were obtained from n=4 mice per condition. **(e)** As in **(a)**, percentage of GFAP-positive staining area was measured. Mean±SD are shown (n= 3-4 for each condition). Representative images of the DG-CA1 region stained with an anti-GFAP antibody (red) and Hoechst dye are shown on the bottom.

**Table S1. Genes and proteins associated with healthy aging.** mRNA (|logFC|>0.5; FDR<5%) and proteins (|logFC|>0.25; FDR<5%) significantly up- or down-regulated when comparing 3- vs. 18-mo. *App^wt^* mice are listed. For each gene, when available, mRNA and protein logFC and FDR are provided. Genes simultaneously found significant at protein and mRNA level are highlighted in gray.

**Table S2. Genes and proteins specifically associated with AD pathology in aged App^NL-F^ mice.** mRNA (absolute logFC>0.5; FDR<5%) and proteins (absolute logFC>0.25; FDR<5%) significantly up- or down-regulated when comparing 3- vs. 18-mo. *App^NL-F^* mice but not found significantly changed in *App^wt^* ageing (see Table S1) are listed. For each gene, when available, mRNA and protein logFC and FDR are provided. Genes simultaneously found significant at protein and mRNA level are highlighted in gray.

**Table S3. Genes and proteins specifically associated with 3xTg-AD pathology.** mRNA (absolute logFC>0.5; FDR<5%) and proteins (absolute logFC>0.25; FDR<5%) significantly up- or down-regulated when comparing 3- vs. 15-mo. 3xTg-AD mice are listed (see Fig. S2c). Absolute logFC and adjusted p-values for 3- vs. 9-mo. *App^NL-G-F^* comparison and the 3- vs. 15- mo. wild-type comparison are also shown.

**Table S4. AD-signatures.** Absolute logFC, adjusted p-values and FDR for mRNA levels of genes identified in the AD-UP and AD-DW signatures. Columns AD-risk, DAM and PIGs indicate whether these genes are included in any given group of genes (*True*, they belong; *False*, they do not). When available, values for the quantified protein are also provided.

**Table S5. Result of the virtual signature-based screening of compounds**. This table shows the detailed scores of the signature-based prioritization of compounds. Compounds selected in at least one of the five queries (Q1-Q5) are shown (8,250 in total). Compounds are identified by InChIKey (ik) and an internal identifier (id); ik is used to rank compounds by default. Molecular weight (mw), Lipinski’s Rule-of-5 violations (ro5), chemical beauty (qed) are given for every compound. In addition, the source of the compound is specified (1/0); alzf: available in AlzForum (i.e., a drug with previously tested against AD), drug: available in DrugBank, pres: part of the Prestwick library, and irb: part of an IRB Barcelona proprietary library. In blue (1/0), we highlight the query (q1-q5) where the compound was a hit; please note that a 0-value in this column does not mean that the compound is not relevant to that query but rather that it was not among the top-scoring hits. The following columns correspond to BBBP, BACE and Aβ activity predictions, based on supervised machine learning (bbbp, bace, ab42, ab40, abratio). In addition, we list putative targets of the molecules (targets), together with a probability score assigned with a simple ligand-based similarity-ensemble approach (tm) (*97*); the column targets_agg simply adds these scores, thereby quantifying the potential promiscuity of the molecule. The remaining columns correspond to the signature-based search. They are organised in colors, red denoting similarity searches against known AD drugs, green transcriptional signature matching, and yellow searches based on the interaction or proximity to putative AD targets. Values shown in these columns correspond to -log10 p-values; these p-values were calculated empirically over the full CC universe (>800k molecules). (Red) Similarity to AD drugs is prefixed with sim_ad. For each molecule, similar AD drugs are listed in sim_ad_drugs (separated by | and separating chemical similarity (sim) and cell-based (phenotypic) similarity (ph) inside the parenthesis); an aggregated score for the overall similarity to AD drugs is given in sim_ad_drugs_agg. Next, similarities are calculated for different AD drug categories, namely amyloid, cholesterol, cholinergic, inflammation, tau, other_neurotrans[mitters], and other. (Green) Transcriptional signature matching is split in five priorities (p4-p0); AD signatures (sigs) identified in human (p4, prefixed jager (*69*)) and our signatures identified in mice (p3, prefixed sbnb, for Structural Bioinformatics & Network Biology) being the categories with top priority, followed by signatures identified in AD cell models (p2, prefixed sbnb_cells), signatures identified in GEO (p1, prefixed geo) and, finally, signatures related to known AD drugs available from LINCS (p0, prefixed with drug InChIKey; in addition, this lowest-priority column contains signatures of proteomics experiments, which we consider less reliable for matching with the transcriptional profiles of the molecules). Columns pX_sigs list the particular signatures matched, together with a connectivity score (con) and a guilt-by-association score (gba), used to relate molecules (based on similarity) to their closest analogs in the LINCS repository (CC space D1) from where transcriptional signatures are available. Columns pX_sigs_agg give an aggregated score for the signature-based search. (Yellow) Compound interaction with putative AD relevant targets. These columns correspond to targets relevant to AD (extracted from OpenTargets, suffixed rational) or targets with a putative association with AD based on LINCS L1000 shRNA experiments (suffixed sh_lincs and following the pX scheme explained above). The relevance of the target to AD is denoted ad:score, corresponding to the OpenTargets score. Interaction of the small molecule with its target is noted with the ligand-based approach (tm); the target of the molecule and the AD target are related with a proximity (prox) score based on the network-based proximity measures developed for the CC (C3-5 levels).

**Table S6. Transcriptional reversion of AD signatures.** List of genes that comprise the leading edge of the transcriptional reversion of the *AD-UP* and *AD-DW* signatures induced by the different treatments. The z-score of the differential expression analysis of treated and untreated *App^NL-G-F^* mice vs. *App^wt^* is provided for each gene. The gene rank of the treated vs. untreated comparison is provided in bins of 1000 (5 = top 1000, 4 = top 2000, 3 = top 3000, 2 = top 4000, 1 = top 5000). Negative signs denote down-regulation. Additional annotations highlight those genes that have been previously related to Alzheimer’s disease or that are up-regulated in disease associated microglia or reactive astrocytosis.

**Table S7.**
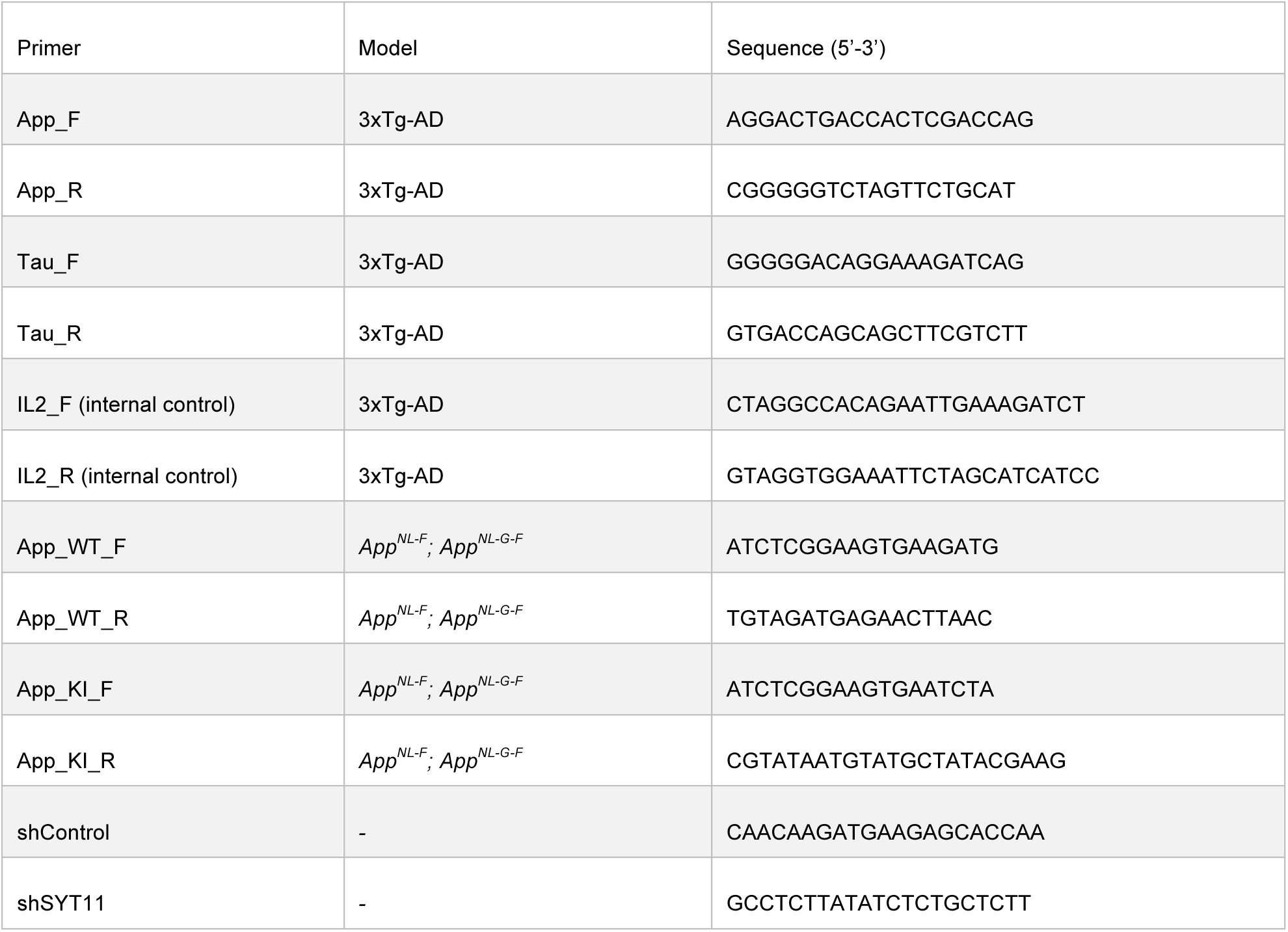
Sequences for mouse genotyping by PCR and shRNA target sequence.

